# Activity-Dependent Alternative Splicing of Adhesion-GPCR Latrophilin-3 Controls Synapse Formation

**DOI:** 10.1101/2023.10.02.560463

**Authors:** Shuai Wang, Chelsea DeLeon, Bryan Roth, Thomas C. Südhof

## Abstract

How synapses are assembled and specified in brain is incompletely understood. Latrophilin- 3, a postsynaptic adhesion-GPCR, mediates Schaffer-collateral synapse formation in the hippocampus but the mechanisms involved remained unclear. Here we show that Latrophilin-3 organizes synapses by a convergent dual-pathway mechanism by which Latrophilin-3 simultaneously activates G_αS_/cAMP-signaling and recruits phase-separated postsynaptic protein scaffolds. We found that cell type-specific alternative splicing of Latrophilin-3 controls its G protein coupling mode, resulting in Latrophilin-3 variants that predominantly signal via G_αs_ and cAMP or via G_α12/13_. A CRISPR-mediated genetic switch of Latrophilin-3 alternative splicing from a G_αS_- to a G_α12/13_-coupled mode impaired synaptic connectivity similar to the overall deletion of Latrophilin-3, suggesting that G_αS_/cAMP- signaling by Latrophilin-3 splice variants mediates synapse formation. Moreover, G_αS_- but not G_α12/13_-coupled splice variants of Latrophilin-3 recruit phase-transitioned postsynaptic protein scaffolds that are clustered by binding of presynaptic Latrophilin-3 ligands. Strikingly, neuronal activity promotes alternative splicing of the synaptogenic variant of Latrophilin-3, thereby enhancing synaptic connectivity. Together, these data suggest that activity-dependent alternative splicing of a key synaptic adhesion molecule controls synapse formation by parallel activation of two convergent pathways, G_αS_/cAMP signaling and the phase separation of postsynaptic protein scaffolds.

---

Synapse formation is central to the assembly of neural circuits in brain. Synapse formation is controlled, at least in part, by trans-synaptic complexes between adhesion molecules that function as signaling modules in organizing pre- and postsynaptic specializations^1,2,3,4^. Among various synaptic adhesion molecules, Latrophilin-3 (Lphn3; gene symbol *Adgrl3*) plays a prominent role in establishing Schaffer-collateral synapses formed by CA3-region axons on CA1-region pyramidal neurons in the hippocampus^5^. Lphn3 belongs to a family of postsynaptic adhesion-GPCRs (aGPCRs) that bind to the presynaptic adhesion molecules teneurin and FLRT^6,7,8^. Lphn3’s function in synapse formation is known to require both its extracellular FLRT/teneurin-binding sequences and its intracellular regions, including its G protein-binding sequence^5,9^, but the molecular mechanisms by which Lphn3 induces synapse formation remains elusive. In cell-signaling assays multiple G_α_ proteins were reported to couple to Lphn3^10,11,12^ and their binding was confirmed by cryo-EM structures^13,14^. However, it is unknown which G_α_ protein physiologically mediates Lphn3- dependent synapse formation, it is also unclear whether G_α_ protein signaling on its own constitutes the core mechanism of synapse formation, and it remains a puzzle how presynaptic ligand-binding to postsynaptic Lphn3 induces synapse formation.

Here, we show that Lphn3 gene (*Adgrl3*) transcripts undergo extensive alternative splicing. The resulting protein variants couple to different G_α_ proteins. Of these, the variant coupling to G_αS_ that induces cAMP production being the predominant splice variant expressed in brain, and is essential for synapse formation in the hippocampus. Moreover, neuronal activity promotes alternative splicing of Lphn3 to induce a shift towards the synapse-forming G_αS_-coupled variant. Strikingly, only the G_αS_-coupled Lphn3 splice variant recruits postsynaptic scaffold proteins by integrating onto the surface of phase-transitioned postsynaptic protein scaffolds. Finally, presynaptic ligands Teneurin and FLRT synergistically promote clustering of Lphn3-containing postsynaptic scaffold protein condensate, explaining how trans-synaptic interaction could induce assembly of postsynaptic specializations. Our data outline a mechanistic pathway of synapse formation in which convergent processes of G_αS_ signaling, phase separation, and trans-synaptic ligand binding mediate the assembly of postsynaptic specializations. This pathway is controlled by the alternative splicing of Lphn3, enabling precise regulation of synapse formation by neuronal activity.

## Extensive alternative splicing of Lphn3 (*Adgrl3*) transcripts

To comprehensively profile the alternative splicing pattern of Lphn3, we analyzed full-length mRNA transcripts from mouse retina and cortex^15^. We identified 5 principal sites of Lphn3 alternative splicing (Figure 1a; Extended Data Figure 1a). Among these, alternatively spliced Exons 6 and 9 encode extracellular sequences that are known to regulate binding to the presynaptic ligand Teneurin^8,16^, while Exon 15 encodes a 13 amino acid sequence within the extracellular GAIN domain. On the cytoplasmic side, Exon 24 encodes a sequence in the 3rd intracellular loop of the 7-TMR GPCR region of Lphn3. The most extensive alternative splicing of Lphn3 is observed in the C-terminal sequence, which is encoded by Exons 28-32. Following the constitutively included Exon 27, all Lphn3 transcripts contain either Exon 31 or Exon 32. Exons 28-30 are variably included in transcripts containing Exon 31 but not in transcripts containing Exon 32. Of these, Exons 28 and 29 are in-frame but Exon 30 can be included in Lphn3 transcripts as Exon 30a (using an internal splice-donor site) or Exon 30b (passing the internal splice-donor site), of which Exon 30b shifts the reading frame of Exon31 (Figure 1b, Extended Data Figure 1b). As a result, Lphn3 transcripts encode three distinct C-terminal sequences. The longest Lphn3 C-terminus is encoded by Exon 31 without or with inclusion of Exons 28, 29, and 30a. The other two shorter C-terminal sequences are either encoded by Exon 32 alone or by the Exon 30b-31 sequence without or with inclusion of Exon 28 and 29 (Figure 1a). Transcriptome analyses of Lphn1 (*Adgrl1*) and Lphn2 (*Adgrl2*) revealed that of the Lphn3 sites of alternative splicing, the sites corresponding to Exons 9 of Lphn3 are conserved in Lphn1 and Lphn2, and that all latrophilins have multiple alternatively spliced 3’ exons, but that the sites corresponding to Exons 15 and 24 of Lphn3 are conserved only in Lphn2 (Extended Data Figure 2a-d).

**Figure 1:**
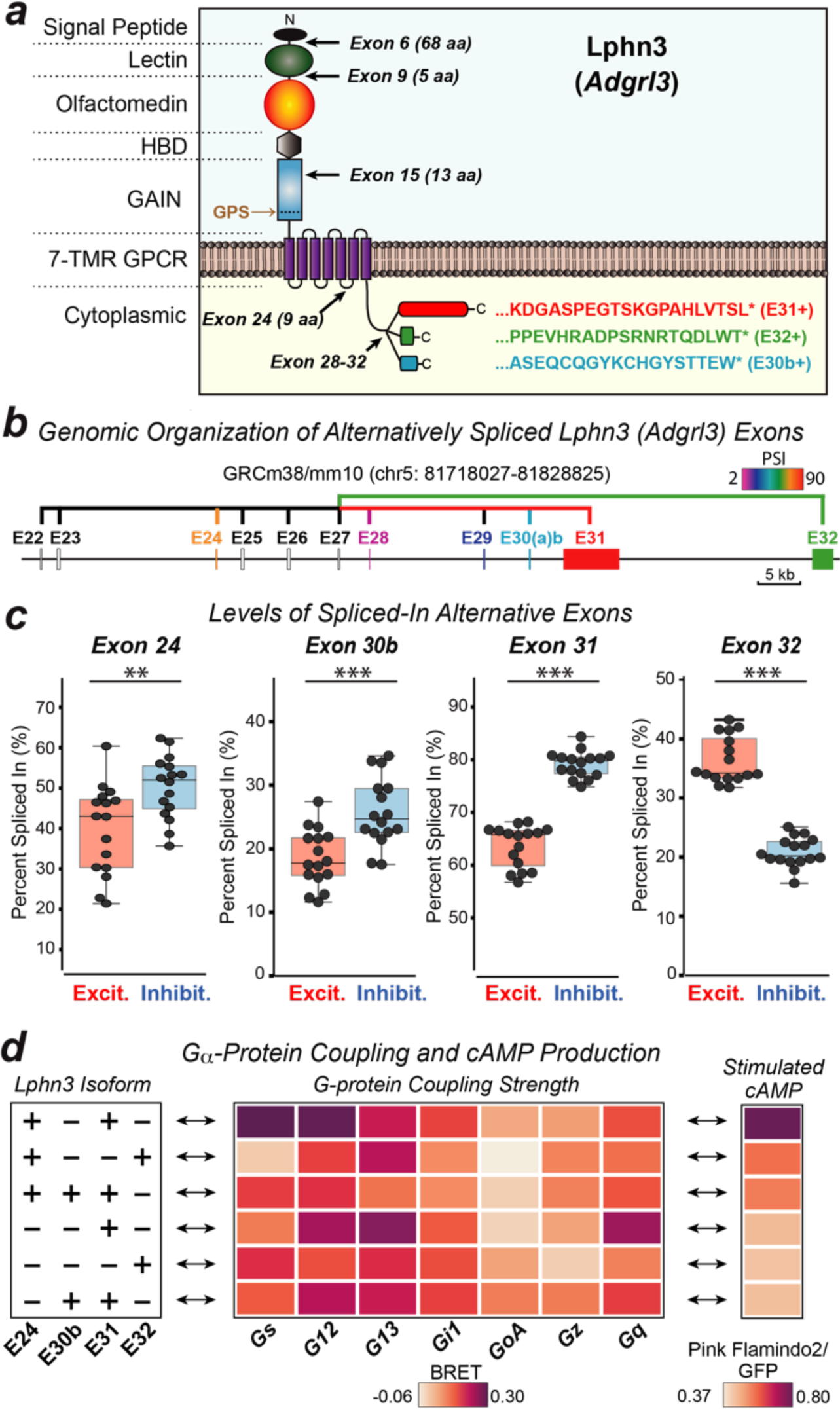
Differentially expressed Lphn3 (*Adgrl3*) splice variants couple to different G- proteins. **a**, Schematic of Lphn3 alternative splicing. **b**, Genomic organization of the 3’ alternatively spliced exons of the Lphn3 gene. Alternative exons are color-coded based on percent spliced-in (PSI) in hippocampus (ED Fig 1c), with constitutive exons colored gray. **c**, Cell type-specific splicing of Lphn3. Raw data from ribosome-associated transcriptome analyses^17^ were analyzed to calculate the percent spliced-in (PSI) of each exon for excitatory and inhibitory neurons (for subtype-specific data, see ED Figure 1c). Each datapoint represents one sample (n=16). Two-sided Student’s t-test was used to calculate the statistical significance (**p < 0.01; ***p < 0.001). **d**, G-protein coupling and stimulated cAMP levels associated with Lphn3 splice variants (left, splice variants; middle, G-protein coupling signal (BRET signal) from TRUPATH assays; right, cAMP stimulated by Lphn3 splice variant expression in HEK293 cells). For detailed data, see ED Figure 4.

To assess the cell-type specificity and relative abundance of various Lphn3 transcripts, we analyzed deep RNAseq data obtained using ribosome-bound mRNAs that were isolated from different types of neurons^17^. We found that mRNAs containing Exon 31 were more abundant (60-80% total) than mRNAs containing Exon 32 (20-40%), with fewer mRNAs containing Exon 30b (20-25%) (Figure 1c; Extended Data Figure 1c). Alternative splicing was cell type-specific, such that inhibitory neurons harbored a higher prevalence of mRNAs containing Exon 31 and 30b than excitatory neurons (Figure 1c), with different subtypes of neurons exhibiting diverse patterns of alternative splicing (Extended Data Figure 1c). Moreover, for some sites of alternative splicing developmental regulation was observed, such as for Exon 30b that was preferentially included early postnatally but excluded during maturation of the brain (Extended Data Figure 3).

## Lphn3 alternative splicing controls G-protein coupling

Given that the alternatively spliced Lphn3 sequences at the cytoplasmic sides are proximal to its G protein interaction site^13,14^, we next systematically analyzed G protein coupling of the principal Lphn3 splice variants (Figure 1d, left) using the TRUPATH resource which affords an unbiased analysis of transducer coupling to GPCRs^18^. The alternative splicing pattern of Exons 24, 30b, 31, and 32 produces 6 principal splice variants that we analyzed. Dramatic differences emerged between Lphn3 splice variants in their G protein coupling preferences (Figure 1d, middle). The most abundant Lphn3 splice variant in the hippocampus (E24+/E30b-/E31+/E32-; Extended Data Figure 1, 3) preferentially couples to G_αS_ and less strongly to G_α12/13_. If Exon 31 is replaced by Exon 32 (E24+/E30b-/E31-/E32+), Lphn3 predominantly couples to G_α12/13_. Inclusion of Exon 30b, or exclusion of Exon 24 also shifts Lphn3 Gα coupling from G_αS_ to G_α12/13_ (Figure 1d; Extended Data Figure 4a). The role of Exon 24 in the 3rd cytoplasmic loop of Lphn3 is consistent with recent studies revealing the importance of this sequence in controlling G protein coupling^19^, but the effect of the C-terminal alternative splicing of Lphn3 on G_α_-protein coupling is surprising given that the sequences involved start 81 residues downstream of the last transmembrane region. These C-terminal sequences are not resolved in present cryo-EM structures of Lphn3 complexed to G proteins^13,14^.

As an orthogonal approach to confirm the TRUPATH data, we measured the ability of various Lphn3 splice variants to increase cAMP levels in HEK293 cells. Upon co-expression of the cAMP reporter Pink Flamindo2^22^ with Lphn3 splice variants in HEK293 cells we observed an increased cAMP level only when the Lphn3 splice variant that preferentially couples to G_αS_ was present (Figure 1d; Extended Data Figure 4b, c). The Lphn3-induced signal was quenched by co-expressed PDE7b, a cAMP-phosphodiesterase^23^, confirming its specificity. We conclude that alternative splicing of Lphn3 controls its G_α_ specificity, with the most abundantly expressed Lphn3 splice variants in the hippocampus preferentially coupling to G_αS_.

Lphn1 and Lphn2 have also been associated with different G_α_ proteins in previous studies ^20,21,9^, prompting us to study their G_α_ protein coupling modes. For Lphn1 and Lphn2 we again observed preferential coupling to G_αS_ for the tested splice variants (Extended Data Figure 2). Viewed together, these data reveal that alternative splicing regulates G_α_ protein preference with G_αS_ being the predominantly coupled G_α_ protein for more abundant latrophilin transcripts, suggesting a possible explanation for the discrepancies among previous studies regarding the signaling pathways controlled by latrophilins.

## CRISPR-mediated genetic manipulation of Lphn3 alternative splicing

To understand whether various Lphn3 splice variants are active in synapse formation under physiological conditions, we focused on its two most abundantly expressed C-terminal exons in the hippocampus: Exon 31 and 32, which are coupled to different G_α_ protein signaling pathways and are alternatively spliced in a mutually exclusive pattern (Figure 1). To control the expression of these two exons from the endogenous Lphn3 gene, as opposed to using overexpressed Lphn3 splice variants in rescue experiments, we designed an acute CRISPR-Cas9 mediated genetic manipulation. In this approach, we selectively deleted the alternatively spliced Exon 31 by targeting its splice-acceptor sequence with a guide-RNA, using a non-targeting guide-RNA as a control, and compared the loss of Exon 31 to the loss of total Lphn3 proteins by targeting the constitutive N-terminal Exon 7 (Figure 2a).

**Figure 2:**
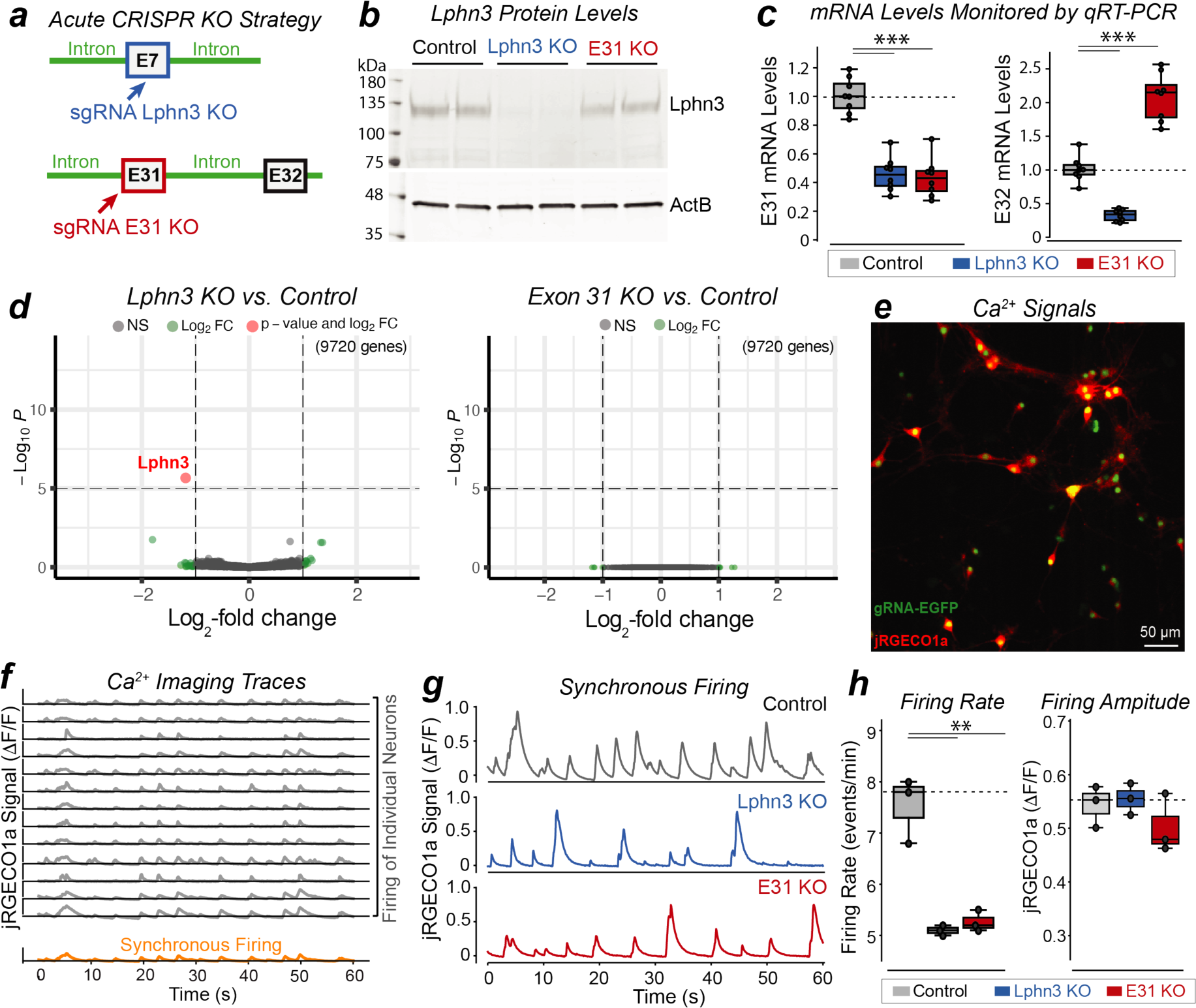
CRISPR-mediated conversion of Lphn3 alternative splicing from Exon 31 to Exon 32 that switches Lphn3 from G_αS_ coupling to G_α12/13_ coupling impairs neuronal network activity. **a**, CRISPR strategy producing either an acute deletion of Lphn3 expression (Lphn3 KO) or a selective deletion of Exon 31 (E31 KO) of Lphn3. **b**, Lphn3 immunoblots showing that the acute Exon 31-specific KO (E31 KO) does not change Lphn3 protein levels, whereas the Lphn3 KO ablates Lphn3 expression. Hippocampal cultures were subjected to CRISPR at DIV3 using lentiviral delivery and analyzed at DIV14; lanes represent independent replicates (control = CRISPR with non-targeting gRNA). **c**, Quantitative RT-PCR (qRT-PCR) measurements demonstrating that the E31 KO and the Lphn3 KO similarly ablate expression of Exon 31-containing Lphn3 mRNAs (left) but have opposite effects on Exon 32-containing Lphn3 mRNAs (right). Exon 31 or Exon 32 junctions with the constitutive Exon 27 were PCR amplified, Ct values were normalized to β actin and all ΔCt values were normalized to controls (CRISPR with non-targeting gRNA). Each data point represents one independent culture (n=8); two-sided Student’s t-test was used to calculate the statistical significance (***p < 0.001). **d**, Gene transcripts with more than 350 reads were used for analyses of differentially expressed genes using DEseq2, shown in volcano plots by comparing Lphn3 KO to control (left), or Lphn3 Exon 31-specific KO to control (right). All data were from 3 independent cultures. Note that only a single gene (Lphn3, the target) has significantly decreased expression by Lphn3 KO, and that no genes were significantly perturbed by Lphn3 Exon31-specific KO. **e**, Representative Ca^2+^-imaging experiment of hippocampal neurons that were infected by lentiviruses expressing a gRNA, EGFP and jRGECO1a^37^. **f**, Representative illustration of the extraction of jRGECO1a signals for individual neurons (gray, top) whose average is the synchronous firing trace for one field of view (orange, bottom). **g**, Representative traces of synchronous firing of neurons analyzed in control, Lphn3 KO and E31 KO neurons for one field of view. **h,** Quantification of synchronous firing rate (left) and amplitude (right) of Ca^2+^-imaging signals. Each datapoint represents one independent culture (n=3); two-sided Student’s t-test was used to calculate the statistical significance (**p < 0.01).

In primary hippocampal cultures, the acute CRISPR-mediated total Lphn3 deletion rendered Lphn3 protein undetectable by immunoblotting, whereas the Exon 31-specific Lphn3 deletion or the control guide-RNA had no apparent effect on Lphn3 protein levels (Figure 2b). Both quantitative RT-PCR and RNAseq analyses from neurons with a targeted deletion of only Exon 31 uncovered a ∼60% decrease in the level of Exon 31-containing mRNAs and a ∼100% increase in the level of Exon 32-containing mRNAs (Figure 2c, Extended Data Figure 5a). When Lphn3 protein was deleted by targeting Exon 7, we observed a ∼60% loss of mRNAs containing Exon 31 or Exon 32, with the decrease in mRNA levels being presumably due to non-sense mediated mRNA decay of the mutant mRNAs^24^.

Transcriptomic analyses detected no off-target effect for either genetic manipulation, and no regulatory function for Lphn3 in gene expression (Figure 2d; Extended Data Figure 5b). These results validate the efficiency and specificity of the CRISPR manipulation approach, with Exon 7 targeting causing a complete loss of Lphn3 protein whereas Exon 31 targeting induces a switch from Exon 31-containing to Exon 32-containing mRNAs.

## Lphn3 coupling to G_α_ protein is essential for synaptic connectivity

We employed three approaches to test whether exclusion of Exon 31, and thus eliminating G_αS_ coupling, affects the function of Lphn3 in synapse formation. First, we measured the network activity of cultured hippocampal neurons using Ca^2+^-imaging (Figure 2e-h). Neurons exhibit regular spiking in culture owing to their spontaneous activity that can be quantified in individual neurons (Figure 2e, f). Averaging the signals of individual neurons in the same recording period produces a “synchronous firing” trace (Figure 2f) that reflects the strength of the synaptic network^25,26^. Quantifications of synchronous firing of cultured neurons showed that the global loss of Lphn3 cause a highly significant decrease (∼40%) in firing rate without altering the signal amplitude (Figure 2g, h). Strikingly, the Exon 31-specific deletion produced a similar decrease in neuronal firing rate as the global loss of Lphn3 proteins (Figure 2g, h).

Second, we asked whether the decrease in firing rate might result, at least in part, from a decrease in excitatory synapse numbers. We quantified the excitatory synapse density in cultured hippocampal neurons after CRISPR-mediated deletions of either all Lphn3 proteins (Exon 7-targeted guide-RNA) or of Lphn3 transcripts containing Exon 31 that couple to G_αS_. Puncta that were positive for both a presynaptic (vGluT1) and a postsynaptic marker (Homer1) were quantified on dendrites (Figure 3a, b). Both the complete loss of Lphn3 and the Exon 31-specific deletion produced a significant decrease in synapse density (Figure 3b).

**Figure 3:**
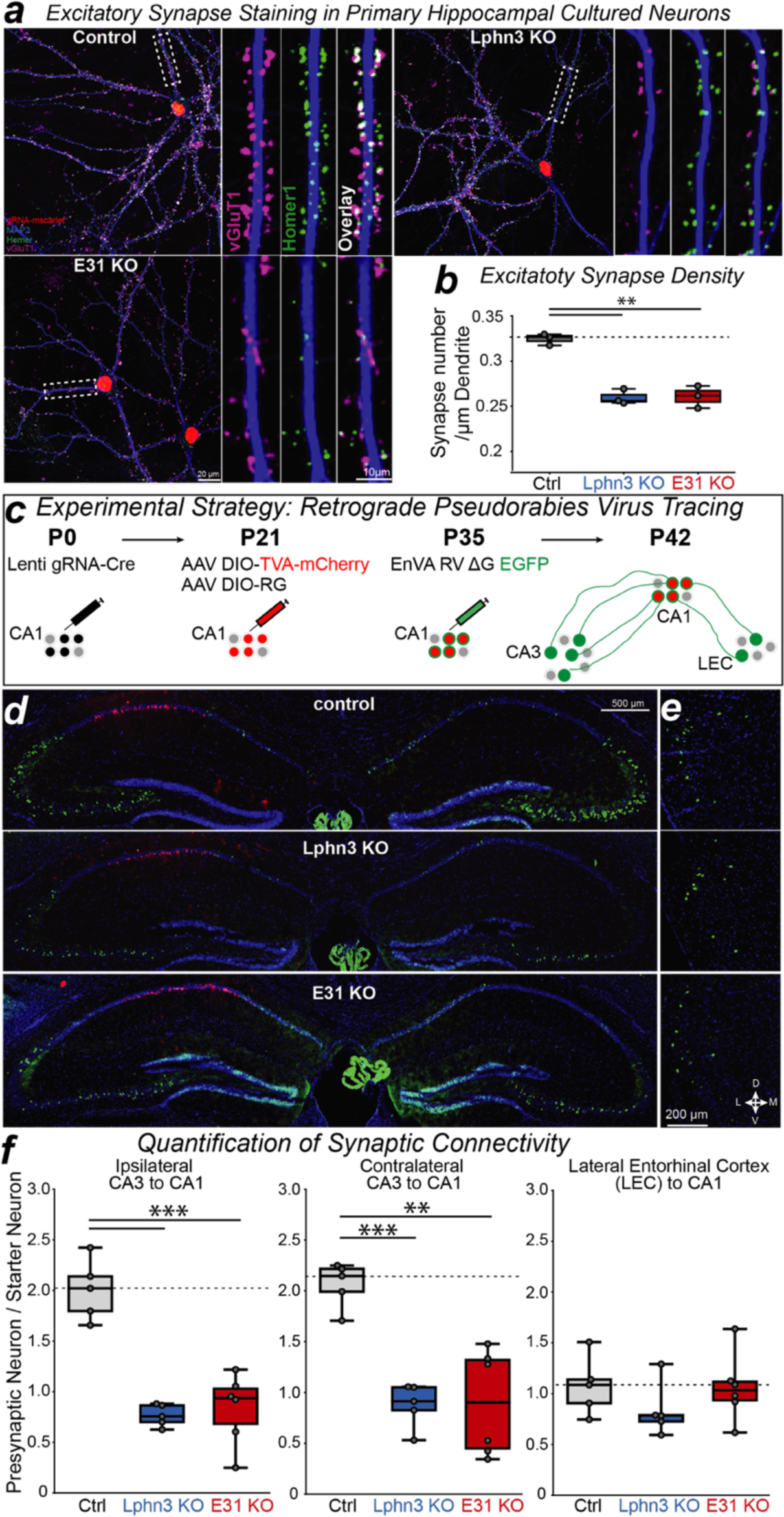
Switching Lphn3 G-protein coupling from G_αS_ to G_α12/13_ by deleting Exon 31 suppresses synaptic connectivity of hippocampal neurons. **a** & **b**, Selective deletion of Lphn3 Exon 31 decreases the excitatory synapse density similarly to the entire deletion of Lphn3 (**a**, representative images of cultured hippocampal neurons stained with antibodies to vGluT1, Homer1 and MAP2; **b**, summary graph of the density of puncta that were positive for both vGluT1 and Homer1). Hippocampal cultures from CAG-Cas9 mice^46^ were transduced with lentiviruses expressing gRNA and nuclear-localized mScarlet-I at DIV3 and analyzed at DIV14. Each datapoint represents an independent culture (n=3); two-sided Student’s t-test was used to calculate statistical significance (**p < 0.01). **c**, Experimental strategy for the retrograde tracing of monosynaptic connections using pseudo-typed rabies virus^39,5^ in CA1 neurons with acute CRISPR-mediated in vivo deletions of Lphn3 or Exon31 of Lphn3. **d** & **e**, Representative images of pseudo-typed rabies tracing experiments in the hippocampal region (**d**) and Lateral Entorhinal Cortex (LEC) (**e**). **f**, Acute CRISPR-mediated in vivo deletion of Exon 31 of Lphn3 (Lphn3 E31 KO) impairs the number of CA3-region input synapses in CA1-region neurons to a similar extent as the overall deletion of Lphn3 (Lphn3 KO) as quantified by retrograde pseudo-typed rabies virus tracing. Box plots show the number of presynaptic neurons (ipsilateral CA3, contralateral CA3, and ipsilateral LEC) normalized to the starter neuron number. Each datapoint represents one animal (n=5-6); two-sided Student’s t-test was used to calculate statistical significance (**p < 0.01; ***p < 0.001). Note that the Lphn3 E31 KO and the Lphn3 KO do not affect entorhinal cortex input synapses which depend on Lphn2 instead of Lphn3^5,47^.

Third, we tested the function of Exon 31 *in vivo*. We used monosynaptic retrograde tracing by pseudo-typed rabies virus to map the connectivity of genetically manipulated starter neurons in the hippocampal CA1 region (Figure 3c). Again, both the loss of all Lphn3 expression and the switching from Exon 31 to Exon 32 caused a large decrease (∼60%) in synaptic inputs to CA1 pyramidal neurons from the ipsi- and the contralateral CA3 region (Figure 3d, f). Inputs from the entorhinal cortex were unchanged since Lphn3 mediates formation of CA3èCA1 Schaffer-collateral but not of entorhinal cortexèCA1 synapses (Figure 3e, f)^5^. These data demonstrate that the Exon 31-containing Lphn3 isoform coupled to G_αS_ is essential for Lphn3-mediated synaptic connectivity.

## Exon 31- but not Exon 32-containing Lphn3 integrates into phase-separated post- synaptic scaffold**s**

Given the importance of the C-terminal sequence of Lphn3 encoded by Exon 31 in synapse formation, we asked whether Exon 31 performs additional functions other than regulating G_α_ protein coupling. We noticed that only the Exon 31-containing transcripts encode the PDZ domain-binding motif at the C-terminus that interacts with Shank proteins^27,28^ (Figure 1a). Thus we sought to test biochemically if full-length Lphn3 could form a complex with postsynaptic scaffold protein networks that are composed of GKAP, Homer, PSD95 and Shank, which are known to form phase-separated protein assemblies^30^ (Figure 4a).

**Figure 4:**
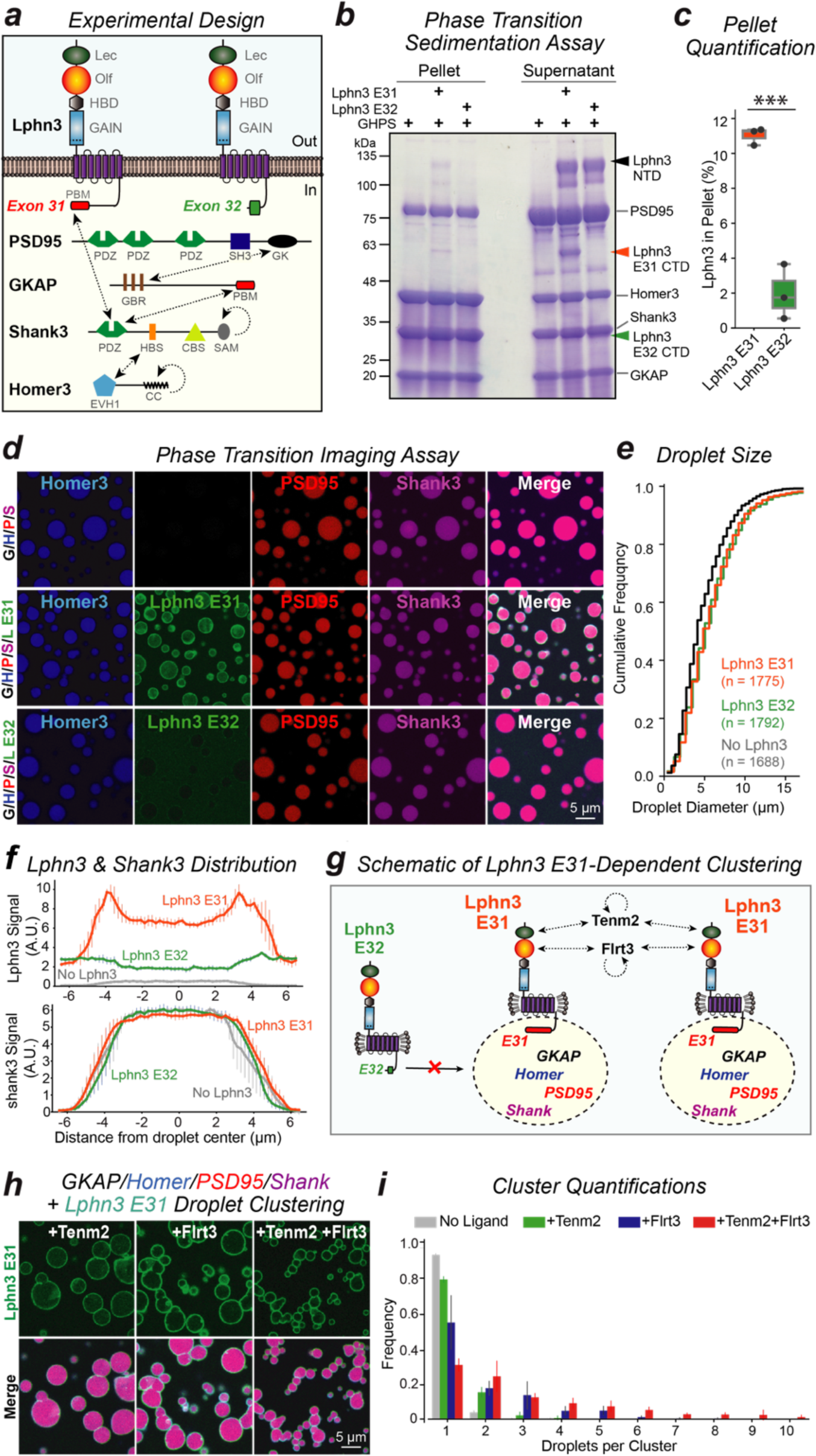
The alternatively spliced Lphn3 variant containing Exon 31 but not Exon 32 spontaneously incorporates into phase-separated assemblies formed by postsynaptic scaffold proteins. **a**, Schematic of proteins (dashed arrows, interactions; circling arrows, homo-oligomerizations; Lec: lectin-like domain, Olf: olfactomedin-like domain, HBD: hormone-binding domain, PBM: PDZ-binding motif, GK: guanylate kinase domain, GBR: GK binding repeat, HBS: Homer-binding sequence, CBS: cortactin-binding sequence, CC: coiled-coil). **b**, Sedimentation assay of phase transition complexes. The scaffold protein mixture containing GKAP, Homer3, PSD95, and Shank3 (GHPS) was incubated with indicated Lphn3 proteins in detergent. Pellet and supernatant were separated by centrifugation and analyzed by SDS-PAGE. Lphn3 is autocleaved at GPS site to produce N-terminal (NTD, black arrow) and C-terminal domains (CTD, red and green arrow)^48^ (see ED Figure 6e, 6f, 7a). **c**, Quantification of Lphn3 pelleting in the sedimentation assay. The bands corresponding to the Lphn3 NTD were used for analysis. Each datapoint represents one independent experiment (n=3); two-sided Student’s t-test was used to calculate the statistical significance (***p < 0.001). **d**, Imaging of phase transitioned complexes. Homer3 (H), Lphn3 (L E31 and L E32), PSD95 (P), and Shank3 (S) were labeled by NHS-ester fluorophore 405, 488, 546, 647, respectively, while GKAP (G) was unlabeled. **e**, Quantification of the sizes of phase-transitioned droplets formed by postsynaptic scaffold proteins GKAP/Homer3/PSD95/Shank3 (G/H/P/S) in the absence of Lphn3 or in the presence of Lphn3 E31 or Lphn3 E32. **f**, Quantification of the Lphn3 (top) and Shank3 fluorescence signal (bottom) across the phase-separated droplet illustrating the surface localization of Lphn3 E31 on the droplet filled with Shank3. Data are means ± SEM (n = 3 independent experiments). **g**, Schematic of the localization of Lphn3 E31 but not of Lphn3 E32 on the surface of phase-transitioned droplets formed by post-synaptic scaffold proteins (GHPS), and the clustering of droplets by Lphn3 ligands Tenm2 and Flrt3. **h**, Representative images of phase-transitioned postsynaptic scaffold protein complexes containing Lphn3 E31 that were clustered by presynaptic ligands Tenm2 and Flrt3 (for further images, see ED Figure 8). **i**, Presynaptic Tenm2 and Flrt3 ligands cluster phase-transitioned postsynaptic scaffolding protein complexes with additive effects of Tenm2 and Flrt3. Data are means ± SEM (n = 3 independent experiments).

Recombinantly purified GKAP, Homer3, PSD95 and Shank3 (Extended Data Figure 6) formed a putative postsynaptic density complex (the “GHPS” complex) via phase separation^30^ that was detected as a sedimented pellet by centrifugation (Figure 4b) and as droplet-like structures by imaging (Figure 4d). When we mixed the GHPS complex with purified detergent-solubilized Lphn3, Lphn3 containing Exon 31 co-sedimented with the GHPS complex, whereas Lphn3 containing Exon 32 largely remained in the supernatant (Figure 4b, c). Moreover, only Exon 31-containing Lphn3 was enriched in the GHPS complex droplet, whereas Exon 32-containing Lphn3 was excluded (Figure 4d-f). Interestingly, the Exon 31-containing Lphn3 was highly enriched in the periphery of the droplets, suggesting that detergent-solubilized Lphn3 formed a layer on top of the post-synaptic scaffold network (Figure 4d, f). These data suggest that alternative splicing of Lphn3 at the C-terminus determines its ability to recruit postsynaptic scaffold proteins, presumably via interactions between the PDZ-binding motif in Exon 31 of Lphn3 and the PDZ domain in Shank3 or/and PSD95.

Teneurins (Tenms)^6^ and Flrts^7^ are single transmembrane region-containing adhesion molecules which bind to the extracellular region of Lphn3. Their binding to Lphn3 is thought to mediate the trans-synaptic interaction between axon terminals and post-synaptic spines^5^. We asked how Tenm2 and Flrt3 might affect the morphology of the above Exon 31- containing Lphn3 in complex with GHPS phase separation condensate. When the purified extracellular region of Tenm2 was added to above complex, we observed a partial clustering of monomeric droplets into dimers (Figure 4g-i). The addition of Flrt3 clustered the droplets into higher-order oligomers, presumably partially due to the higher affinity of Flrt3 (*K_d_* = ∼15 nM)^7^ than Tenm2 (*K_d_* = ∼500 nM)^31^ for Lphn3. Tenm2 and Flrt3 can bind to Lphn3 simultaneously^31,32^ and acted synergistically in promoting the clustering of phase-transitioned droplets (Figure 4g-i). The clustering effect was not observed in Exon 32- containing Lphn3 (Extended Data Figure 8). Because Tenm2 is an obligatory dimer via disulfide bonds between EGF repeat domains^33^ and Flrt3 forms dimers via its leucine-rich repeat domain^34,35^, we posit that the dimerization of ligands promoted the intermolecular interaction of Lphn3 in adjacent droplets, resulting in the formation of clustered Lphn3- coated postsynaptic scaffold protein condensates (Figure 4g). Together, these data suggest Exon 31 of Lphn3 recruits postsynaptic scaffold proteins to form the protein-dense compartments, which are assembled into higher-order clusters in the presence of pre- synaptic ligands (Teneurins and FLRT).

## Neuronal activity promotes the splicing of synapse-forming variant of Lphn3

Given the observed cell type-specific expression of Lphn3 alternative splice variants and their distinct functions in synapse formation, we wondered whether alternative splicing of Lphn3 is regulated by neuronal activity. To this end, we analyzed a recent transcriptomic dataset in which the neurons in culture or in vivo were examined after stimulation^36^. No activity-dependent changes in the inclusion of Exon 24 of Lphn3 mRNAs were found, whereas a significant shift of alternative splicing in Exons 31 and 32 was detected (Figure 5a, b). Specifically, both depolarization-induced stimulation in cultured neurons (Figure 5a) and kainate-induced activation of hippocampus in vivo (Figure 5b) caused a shift from Exon 32-containing to Exon 31-containing Lphn3 mRNAs, with the abundance of Exon 32- containing mRNAs almost halved after stimulation. Thus, neuronal excitation shifted Lphn3 towards the alternative splice variant that promotes synapse formation.

**Figure 5:**
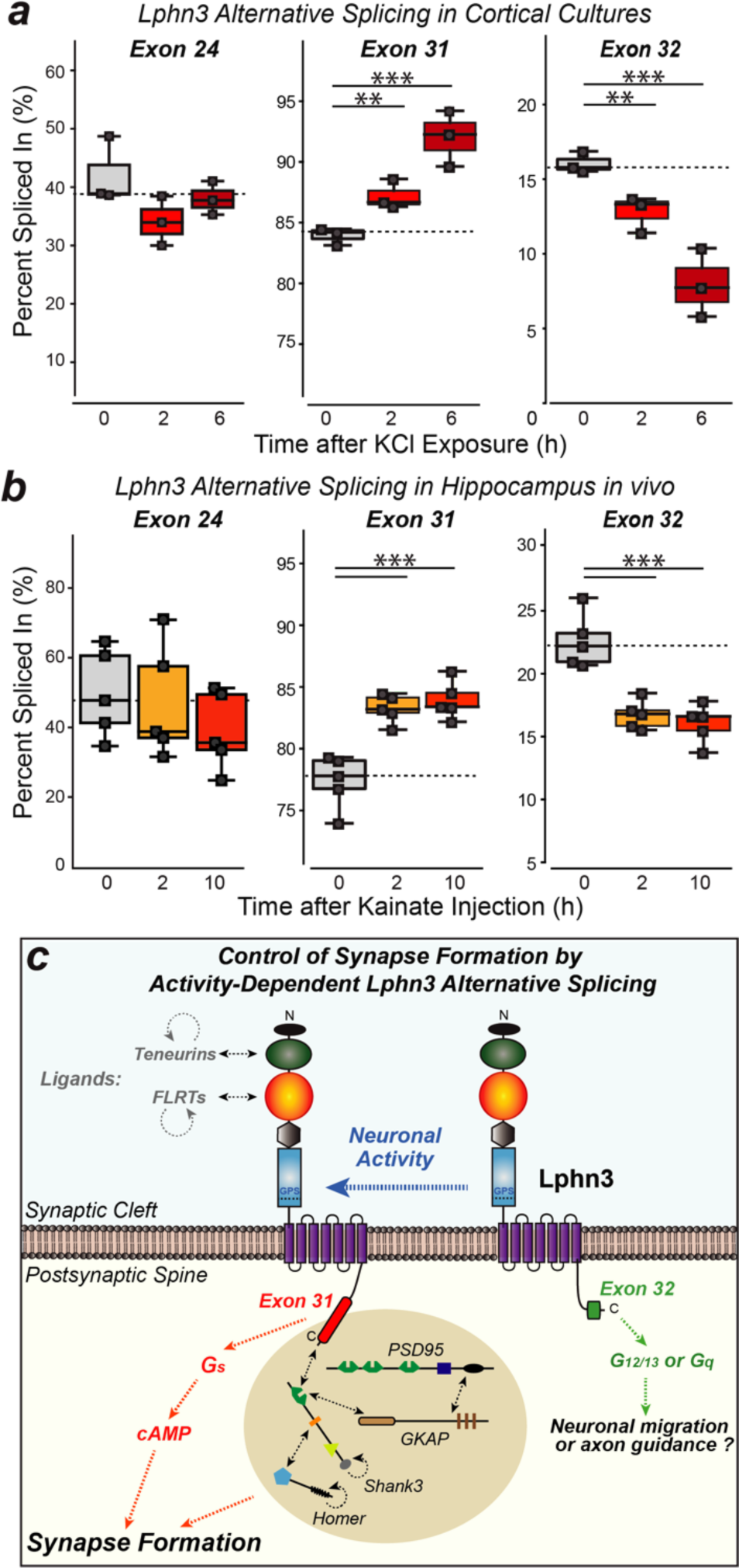
Neuronal activity regulates Lphn3 alternative splicing, leading to an increase in Exon 31 inclusion that promotes synapse formation. **a** & **b**, Neuronal activity induced by KCl depolarization in cortical cultures (**a**) or by kainate injections in vivo (**b**) increases the inclusion of Exon31 and decreases the inclusion of Exon 32 in Lphn3 mRNAs without affecting the splicing of Exon 24. Raw data is analyzed from previous study^36^. Each datapoint represents one replicate (n=3-5). A two-sided Student’s t-test was used to calculate statistical significance (*p < 0.05; **p < 0.01; ***p < 0.001). **c**, Model of the mechanism of action of Lphn3 in synapse formation and the regulation of Lphn3 function by alternative splicing of Exon 31.

## Summary

Here we show that Lphn3 transcripts are subject to extensive alternative splicing that control its G protein coupling specificity and the recruitment of postsynaptic scaffolding. We demonstrate that the Lphn3 splice variant which is coupled to G_αS_ and mediates cAMP signaling is required for synapse formation in vivo, and that this splice variant selectively recruits postsynaptic scaffolding by enabling the incorporation of Lphn3 onto the surface of phase-transitioned postsynaptic density protein complexes, which further clustered into larger assemblies by interacting with presynaptic ligands. These findings outline a synapse formation mechanism that is mediated by two parallel but convergent pathways, localized G_αS_/cAMP signaling and phase transitioning of postsynaptic scaffolding proteins which are orchestrated by Lphn3.

## Acknowledgements

We thank J.H. Trotter and Z.J. Sun for help on imaging; K. Liakath-Ali, X.D. Chen and J.Y. Dai for helpful discussions. This work was supported by grants from National Institute of Mental Health to T.C.S (5R01 MH126929-02) and a Stanford Maternal & Child Health Research Institute Postdoctoral Support grant to S.W. (1220319-117-JHACT), and R24DK116195, the NIMH Psychoactive Drug Screening Program and the Michael Hooker Distinguished Professorship to B.L.R.

## Author Contributions

S.W. performed all experiments and analyzed all data, except the TRUPATH assay which was performed by C.D. and supervised by B.L.R.. S.W. and T.C.S. conceptualized the project, designed the experiments, and wrote the manuscript with input from all authors. All authors contributed to data analyses.

## Data Availability

The data supporting the findings of this study are deposited in the Stanford Data Repository (https://purl.stanford.edu/nj297xj2116) except for the RNAseq data that will be deposited at Gene Expression Omnibus repository with accession code GSEXXXXXXX.

## Code Availability Statement

The codes used for this study will be deposited in the Stanford Data Repository (https://purl.stanford.edu/nj297xj2116). Further details are available from the corresponding author upon request.

## Method

### Mouse handling

C57BL/6 (JAX # 000664) mice were used for tissue RT-PCR experiment, and CAG- Cas9 mice (JAX #024858) were used for all other experiments. Mice were weaned at postnatal day 21 and housed in groups of maximum 5 on a 12 hr light/dark cycle with food and water *ad libidum*, in Stanford veterinary service center. All procedures conformed to National Institutes of Health Guidelines for the Care and Use of Laboratory Mice and were approved by the Stanford University Administrative Panel on Laboratory Animal Care.

### Plasmids

Plasmids for the TRUPATH assay were from the Roth lab (Addgene #1000000163). Pink flamindo 2, GFP, and PDE7b constructs were from previous study^9^. Mouse Lphn3 of specified splicing variants (all have the following splicing configuration: -E6/+E9/+E15/-E28/-E29) were cloned into pCMV vector using In-Fusion® HD assembly. For manipulating Lphn3 KO and Lphn3 E31-only KO, gRNAs were cloned into lentiCRISPR v2 (Addgene #52961), followed by hsyn1 promoter driven EGFP (for calcium imaging and RNAseq), or mScarlet-I (for synapse puncta staining), or Cre recombinase (for monosynaptic rabies tracing), using In-Fusion® HD assembly. jRGECO 1a^37^ was cloned into FSW lentiviral vector. GKAP, Homer3, PSD95, Shank3 coding regions^30^ containing N-terminal 6xHis and 3C protease cleavage site were cloned into pCT10 vector. N-terminal flag tagged- Lphn3 E31 and Lphn3 E32 (with full splicing combination: E31: -E6/+E9/+E15/+E24/-E28/-E29/- E30/+E31, E32: -E6/+E9/+E15/+E24/-E28/-E29/-E30/+E32) were cloned into lenti_2nd_CMV- TetO2_IRES-GFP vector (Addgene #113884) while removing IRES-GFP. Sequences of all constructs were confirmed by Sanger sequencing at Elim Biopharm Inc, or long-read sequencing at Primordium.

### Genetic CRISPR manipulations

3 gRNAs were designed in this study. The control gRNA was designed to have no target in the mouse genome. The Lphn3 KO gRNA targets the constitutive exon 7 to induce frameshift. The Lphn3 E31 gRNA targets the splicing acceptor site immediate upstream of exon31, to disrupt the inclusion of exon 31. Potential off-targets were assessed by Cas-OFFinder^38^ to ensure specificity.

### Generation of reference exon list

The exon coordinates of Lphn3 were extracted from GFF annotation of mouse genome GRCm38/mm10. Non-overlapping exons were named numerically in ascending order from 5’ to 3’ of the transcript. For exons with overlapping region (mostly due to alternative splicing donor/acceptor site), they were named with the same number but different alphabets. For exons at the 5’ and 3’ UTR only the longest annotated exons were used, since current study focuses on the coding region. This generated the draft of exon list. Because annotated exon list may contain exons which never translate to proteins (mostly due to incomplete splicing / incorrect annotation), the draft exon list was used to map the reads of Lphn3 from Ribotaq sequencing dataset^17^ which is highly enriched for translating mRNAs, during which only exons detected in this dataset was preserved to produce the final reference exon list for this study.

### Analysis of high-throughput sequencing data

This study analyzed three datasets from published studies and one dataset generated from this study:

1. Pacbio long-read mRNA sequencing data^15^ were downloaded from NCBI BioProject repository (accession number PRJNA547800). Fastq files were aligned to reference genome (GRCm38/mm10) using gmaps. Reads belonging to Lphn3 from above 5 tissue samples (4 developmental stages of retina and P35 of cortex) were combined to increase the read depth, for analyzing the abundance of full-length transcripts.
2. Cell type-specific Ribotaq sequencing data^17^ were downloaded from GEO (accession code: GSE133291). Fastq files were aligned to reference genome (GRCm38/mm10) using STAR. Reads belonging to Lphn3 were used for calculating percent spliced in (PSI) of exons.
3. Neuronal activity regulated transcriptome dataset^36^ were downloaded from GEO with accession number GSE175965. Fastq files of cortical culture stimulated by KCl, and hippocampal tissue from C57BL/6N mice injected with kainic acid were aligned to reference genome (GRCm38/mm10) using STAR. Reads belonging to Lphn3 were used for calculating percent spliced in (PSI) of exons.
4. The fastq files of Lphn3 KO and Lphn3 E31 KO studies of this work were aligned to reference genome (GRCm38/mm10) using STAR. For differential expression gene analysis at transcriptome level, HTseq was used to count reads per gene from alignment files. Only genes with more than 350 reads per sample were retained, resulting in 9750 genes for downstream analysis. DEseq2 was used to calculate the adjusted p value and product volcano plots using default settings. Reads belonging to Lphn3 were used to calculate RPKM of exon31 and 32.

### Calculation of exon percent spliced in (PSI)

PSI is the percentage of reads containing target exon among all reads at target region. For alternative exon containing both 5’ and 3’ flanking exons (exon 24 and 30b for example), only the reads spanning the target exon-exon junctions were used for calculation. For alternative exons at the 3’ termini (exon31 and exon32), all reads containing the target exon were used for calculation since the 3’ termini of Lphn3 ends with either exon 31 or 32. Exon 31 and 32 reads were normalized by exon length before calculating PSI.

### Sample preparation of Lphn3 KO and Lphn3 E31 KO neurons for next-generation sequencing

Primary hippocampal culture neurons were infected with lentiviruses expressing gRNAs (control, Lphn3 KO, Lphn3 E31-only KO) at DIV3 and maintained till DIV14. One coverslip of culture was resuspended with 200 µl TRIzol® on ice, mixed with 50 µl chloroform, and incubated at RT for 2 min. Samples were centrifuged at 12000 g for 15 min at 4°C in Eppendorf 5417C centrifuge. The aqueous layers were added to 100 µl ice-cold isopropanol for thorough mixing before incubation at -80°C for 1 hr. Samples were thawed on ice and centrifuged at 20817 g for 20 min at 4°C in Eppendorf 5417C centrifuge. Pellets were washed by 0.5 ml ice cold 75% EtOH before centrifuged at centrifuged at 20817 g for 10 min at 4°C, and subsequently resuspended by 40 µl ddH2O containing 0.2 U/µl SUPERase•In™ RNase Inhibitor. Total RNA samples were converted to library using Illumina Stranded mRNA kit and sequenced in NovaSeq (PE 150) with 40M paired reads at Medgenome Inc.

### TRUPATH G-protein coupling assay

HEK293T cells were obtained from ATCC and maintained, passaged and transfected in DMEM medium containing 10% FBS, 100 U ml^−1^ penicillin and 100 µg ml^−1^ streptomycin (Gibco-ThermoFisher) in a humidified atmosphere at 37 °C and 5% CO_2_. After transfection, cells were plated in DMEM containing 1% dialyzed FBS, 100 U/mL penicillin, and 100 µg/mL streptomycin for BRET assays. Constitutive activity of Lphns were accomplished using the previously optimized Gα-Rluc8, β-subunit, and N-terminally tagged γ-GFP2 subunit pairs described before^18^. HEK cells were plated in a 12-well plate at a density of 0.3-0.4x10^6^ cells per well with DMEM containing 10% FBS, 100 U/mL penicillin, and 100 µg/mL streptomycin. Six hours later, cells were transfected with a 1:1:1 ratio of optimized Gα:ý:γ pairings at 100 ng and various amounts of receptor (25 ng, 50 ng, 100ng, 200 ng, 300 ng) using TransIt-2020 (Mirius Bio). To establish a baseline for the cells, pcDNA was used at 100 ng and referred to as 0 ng. The following day, cells were removed from the 12-well plate with trypsin and seeded in a 96-well white, clear-bottomed plate (Greiner Bio-One) with DMEM containing 1% dialyzed FBS at a cell density of 30,000-35,000 cells per well. Cells were incubated overnight to allow for attachment and growth. The next day, media was aspirated from the wells. A solution of assay buffer (20 mM HEPES, Hank’s balanced salt solution, pH 7.4) and 5 μM of coelentrazine 400a (Nanolight Technology) was prepared and added to each well. Cells were allowed to equilibrate with the coelentrazine 400a in the dark for 10 minutes. Corresponding BRET data was collected using a Pherastar FSX Microplate Reading with luminescence emission filers of 395 nm (RLuc8-coelentrzine 400a) and 510nm (GFP2) and an integration time of 1 sec per well. BRET ratios were calculated as a ratio of the GFP2:RLuc8 emission. The constitutive coupling (0 ng) was used as the baseline to subtract NET BRET of the experimental conditions for each receptor. 3 independent cultures with 7 technical replicates in each culture were used in total.

### cAMP reporter assay

HEK293T cells were maintained in DMEM+10% FBS at 37°C 5% CO2, and seeded onto 24-well plate. During calcium transfection of each well, EGFP (0.23 µg), pink flamindo2 (0.2 3µg), GαS (0.16 µg), Gβ (0.16 µg), Gγ (0.16 µg) were used for all conditions. When indicated, additional constructs were co-transfected including PDE7b (0.23 µg) and 6 isoforms of Lphn3 (E24+/E30b-/E31+/E32-, E24+/E30b+/E31+/E32-, E24+/E30b-/E31-/E32+, E24-/E30b-/E31+/E32-, E24-/E30b+/E31+/E32-, E24-/E30b-/E31-/E32+, 0.23 µg each). 16 hr post-transfection, medium was replaced by 0.5 ml DMEM+10% FBS. 36-48 hr post transfection, medium of all cultures were replaced by 0.5 ml imaging buffer (20mM Na-Hepes pH7.4, 1x HBSS (gibco #14065056)) and incubated at RT for 30 min. When indicated, 2.5 µM Forskolin and 5 µM IBMX added to the culture for 5 min. Imaging was performed under Nikon confocal microscopy at 10x objective.

### Primary hippocampal neuron culture

Neonatal P0 mice pups of CAG-cas9 mice (JAX #024858) were dissected in ice cold HBS to obtain hippocampi, which were digested in 1% v/v papain suspension (Worthington) and 0.1 U/µl DNaseI (Worthington) DNaseI for 15min at 37°C. Hippocampi from 2 pups were washing with calcium-free 1xHBS (pH7.3) and dissociated using gentle pipetting in plating medium (MEM containing 5% FBS, 0.6% glucose, 2% Gem21 NeuroPlex™ Supplement, 2 mM GlutaMAX™), filtered through 70 µm cell strainer, and seeded onto Corning® Matrigel®- coated 12 mm cover glasses in one 24-well plate, and maintained at 37°C 5% CO2. 16 hr post seeding (DIV1), 90% media were replaced by maintenance medium (Neurobasal A with 2% Gem21 NeuroPlex™ Supplement, 2 mM GlutaMAX™). At DIV 3, 50% of medium were replaced with fresh maintenance medium supplemented 4 µM Ara-C (Cytosine β-D-arabinofuranoside hydrochloride), and lentivirus expressing gRNA. When indicated, lentiviruses expressing jRGECO1a were added at DIV7. At DIV7, 10, and 13, 30% media were replaced with fresh maintenance medium, before analyzing at DIV14.

### Virus preparation

Lentiviruses were produced in HEK293T cells using the 2nd generation packaging system. Per 150 cm^2^ of cells, 186 µl 2 M CaCl_2_ containing 5.8 µg of lentivirus shuttle vector, 2.5 µg pVSVG (addgene 12259), 4.2 µg Gag-Pol-Rev-Tat (addgene 12260) at a total volume of 1.5 ml was added dropwise to an equal volume of 2X-HBS (280 mM NaCl, 10 mM KCl, 1.5 mM Na_2_HPO_4_, 12 mM glucose, and 50 mM HEPES, pH 7.11) under constant mixing, incubated for 15 min at room temperature, and added dropwise to the cells. 8-12hr post-transfection, medium was replaced by DMEM with 10% FBS. 48 hr post-transfection, cell medium was cleared by spinning at table-top centrifuge at 2000 g for 3 min, and filtered via 0.45 µm PES membrane. The viral supernatant was loaded onto 2 ml 30% sucrose cushion in PBS and centrifuged in Thermo Scientific SureSpin 630 rotor at 19000 rpm for 2 hr. Viral pellet was resuspended in 30 µl MEM and flash frozen in liquid nitrogen. AAVs (CAG-DIO-RG and CAG-DIO-TCB-mCherry) in capsid 2.5 and pseudotyped rabies virus RbV-CVS-N2c-deltaG-GFP (EnvA)^39^ were prepared at Janelia Farm Viral core facility.

### Monosynaptic retrograde rabies tracing

P0 neonatal mouse pups were anesthetized on ice for 4 min and head-fixed on by ear bars and 3D-printed mold. 0.35 µl 1x 10^9^ IU/ml lentiviruses (hsyn1-gRNA-NLS-cre) were injected unilaterally to CA1 at coordinates AP +0.95 mm, ML -0.92 mm, DV -1.30 mm (zeroed at Lambda). At P21, mice were anesthetized by avertin (250 mg/kg) and head-fixed on stereotaxic injection rig, 0.2 µl AAVs CAG-DIO-RG 3.6 x 10^12^ GC/ml and CAG-DIO-TCB-mCherry 6.35 x 10^12^ GC/ml (1:1 volume mix) were co-injected to CA1 at coordinates AP -1.80 mm, ML -1.35 mm, DV -1.30 mm (zeroed at Bregma). At P35, the same CA1 site was injected by 0.15 µl EnvA- pseudo-typed rabies virus RbV-CVS-N2c-deltaG-GFP at 2 x 10^8^ IU/ml. After the surgery, the incisions of P21 and P35 mice were closed by suture and 3M Vetbond tissue adhesive (# 1469SB). After all injections, mice were recovered on a heating pad before returning to home cage. At P42, mouse brains which had been perfused were fixed in 4% PFA (Electron Microscopy Sciences, EM grade #15714) in PBS for 4hr at RT, subsequently incubated in 30% w/v sucrose in PBS at 4°C overnight, and cryopreserved in Tissue-Tek® O.C.T. compound (Sakura) on dry ice. Frozen tissue blocks were cut into 20 µm coronal sections on a cryostat and collected on glass slides (Globe Scientific #1358W). Sections were air-dried, stained in 1 µg/ml DAPI for 10 min, washed once with PBS, and sealed in Fluoromount-G® (Southern Biotech, 0100-01). Sections were imaged on Olympus VS200 slide scanner at 10x. 5-6 mice were used per condition for the study.

### RT-PCR of Lphn3 exons in tissues

C57BL/6 Mice at postnatal day (P4, P9, P14, P21, and P35) were euthanized and brains were dissected to isolate olfactory bulb, cerebellum, hippocampus, prefrontal cortex, striatum and retina. Tissues were grinded with 500 µl TRIzol® on ice, mixed with 125 µl chloroform, and incubated at RT for 2 min. Samples were centrifuged at 12000 g for 15 min at 4°C in Eppendorf 5417C centrifuge. The aqueous layers were added to 250 µl ice-cold isopropanol for thorough mixing before incubation on ice for 5 min. Samples centrifuged at 20817 g for 20 min at 4°C in Eppendorf 5417C centrifuge. Pellets were washed by 0.5 ml ice cold 75% EtOH before centrifuged at centrifuged at 20817 g for 10 min at 4°C, and subsequently resuspended by 40 µl ddH2O containing 0.2 U/µl SUPERase•In™ RNase Inhibitor. 100 ng of total RNA was used for cDNA conversion by PrimeScript™ RT-PCR Kit using random 6mer. 1 µl cDNA was used for PCR targeting exon-exon junction regions using Ex Taq™ DNA Polymerase. The following primers were used: β- actin (5’-TCTACAATGAGCTGCGTGT-3’, 5’-CGAAGTCTAGAGCAACATAG-3’), Lphn3 E6 (5’- CCACAGCTACTCATCCTCAC-3’, 5’-GCTCTCGATCATGATGACGT-3’), Lphn3 E15 (5’- GGGGACATCACCTACTCTGT-3’, 5’-TCAGGTCTCTCCAGGCATTC-3’), Lphn3 E24 (5’- CCTGAATCAGGCTGTCTTGA-3’, 5’-AAATGGTGAAGAGATACGCC-3’), Lphn3 E31 (5’- TCCAGGACGGTACTCCACA-3’, 5’-GGCATTGTTCAGAAGCCCCT-3’), Lphn3 E32 (5’- TCCAGGACGGTACTCCACA-3’, 5’-TCCTGTGTCCTGTTTCGGGA-3’). PCR program is: 94° 1min, 94°C 30s, 55°C 30s, 72°C 1min, go to step2 for 30 times. PCR products were separation on 2% agarose gel in 1xTAE buffer and imaged using Biorad Gel imaging system.

### RT-qPCR of Lphn3 KO and Lphn3 E31 KO neurons

80ng total RNA for each culture was used for converting to cDNA by PrimeScript™ RT-PCR Kit using random 6mer. 1µl cDNA was used for qPCR experiment in TaqMan™ Fast Virus 1-Step Master Mix using PrimeTime Std® qPCR designed primer-probe sets (For β-actin: 5’-GACTCATCGTACTCCTGCTTG-3’, 5’- GATTACTGCTCTGGCTCCTAG-3’, /56-FAM/CTGGCCTCA/ZEN/CTGTCCACCTTCC/3IABkFQ/; for Lphn3 E27-E31 junction: 5’-CCTTCATCACCGGAGACATAAA-3’, 5’-GTGGTAGAGTATCCATGACACTTG-3’, /56-FAM/CA GCTCAGC/Zen/ATCGCTCAACAGAGA/3IABkFQ/; for Lphn3 E27-E32 junction: 5’- CAGTCAGAGTCGTCCTTCATC-3’, 5’-GTCAGTCTCAGGTCCATAAGTC-3’, /56- FAM/AACAGCTCA/Zen/GCATCGCTCAACAGA/3IABkFQ/). PCR program is: 95° 20s, 95°C 3s, 60°C 30s, go to step2 for 40 times. Ct values for Lphn3 E31 and E32 sample were subtracted by that of β-actin from the same sample to get ΔCt. All ΔCt values were normalized by control gRNA. 8 cultures were used in total.

### Immunoblotting analyses

One well of neuron culture from 24 well plate was lysed in 50µl lysis buffer (20mM Tris pH7.5, 500mM NaCl, 1% Tritonx100, 0.1% SDS 1xRoche EDTA-free protease inhibitor) at RT for 5min. 20µl 5x SDS loading buffer was added and the samples were subject to SDS-PAGE electrophoresis. Gels were transferred to 0.2µm nitrocellulose membrane in Trans-Blot Turbo Transfer system (Biorad) and blocked by western blocking buffer (5% BSA in 1xTBST) at RT for 30min. Mouse anti-Lphn3 (Santa Cruz Biotech #sc-393576, 1:1000) and mouse anti-actin (Sigma #A1978, 1:3000) antibodies in western blocking buffer were added and incubated at 4°C for overnight. The membranes were washed in western blocking buffer three times for 10min each, and IRDye® 800CW Donkey anti-Mouse IgG Secondary Antibody (Li-cor # 926-32212, 1:20000) in western blocking buffer was added to the membrane which was incubate at RT for 1hr and washed in 1xTBST three times for 10 min each. Samples were imaged at Odyssey Imager (Li-Cor).

### Calcium imaging

Primary culture neurons were maintained as mentioned above, except they were infected with lentivirus expressing hsyn1-gRNA-EGFP at DIV3, and lentivirus expressing hsyn1- jRGECO1a at DIV7. At DIV14, the coverslips containing neuron were washed once with 37°C warmed Tyrode buffer (25mM Na-HEPES pH7.4, 129mM NaCl, 5mM KCl, 2mM CaCl_2_, 1mM MgCl_2_, 15mM glucose and transferred to 12 well glass plate (Cellvis # P12-1.5H-N) in Tyrode buffer. After 30 min of incubation in Tyrode buffer at 37°C 5% CO2, the cultures were imaged in Leica microscopy at 37°C 5% CO2, with 50 ms exposure, 85 ms interval for 1 min for each field of view (FOV). 6-8 FOVs were recorded for each coverglass of culture. For each condition from one batch of culture, 3- 5 cover glasses of cultures were imaged. 3 batches of culture were used in total.

### Immunohistochemistry and synapse puncta imaging

Primary hippocampal neurons were washed in Tyrode buffer (25 mM Na-HEPES pH 7.4, 129 mM NaCl, 5 mM KCl, 2 mM CaCl_2_, 1 mM MgCl_2_, 15 mM glucose) and fixed in 4% PFA, 4% sucrose in 1 x DPBS at 37°C for 15 min. Afterwards, neurons were washed three times with 1xDPBS for 5 min each, and permeabilized in 0.1% Triton X-100 in 1x DPBS for 10min at RT without shaking. After blocking with 0.5% fish skin gelatin in 1x DPBS at 37°C for 1hr, culture was stained with chicken anti-MAP2 (Encor #CPCA-MAP2, 1:1000), guinea pig anti-vGluT1 (Milipore #AB5905, 1:1000), and rabbit anti-Homer (Milipore #ABN37, 1:1000) in blocking buffer at 4°C for overnight. Samples were washed three time with 1xDPBS for 8 min each, and incubated with secondary antibodies (anti-chicken Alexa405, anti-guinea pig Alexa647, and anti- rabbit Alexa488) in blocking buffer at 37°C for 1hr. Afterwards, the culture coverslips were washed three times with 1x DPBS for 8 min each, once with ddH2O briefly, before loaded onto glass slide slides (Globe Scientific #1358W) in Fluoromount-G® (Southern Biotech, 0100-01) and sealed in nail polish (Amazon #B000WQ9VNO). Samples were imaged under Nikon confocal microscopy at 60x, with 0.35 µm step size and 4-6 z-stacks. For each coverglass of culture, 20-25 neurons containing well-isolated dendrites were imaged. For each condition of one batch of culture, 2 cover glasses of culture were imaged. 3 batches of culture were used in total.

### Image analyses

Four types of image analyses were performed.

1. **Quantification of excitatory synapse puncta density.**Maximum intensity files were produced from z-stacked images. Background was subtracted and the 5-10 well-isolated secondary dendrites were cropped from each neuron in Fiji (v 2.9.0) for processing. The cropped files were converted to binary images using the same threshold for the same channel, for the same batch of experiment. For calculating excitatory synapse puncta, the vGluT1 and Homer binary images were used to product overlapped region. For calculating vGluT1 and Homer puncta separately, vGluT1 and Homer binary images were directly used for downstream analysis. Binary regions containing more than 2 neighboring pixels was considered as puncta, which were searched and quantified using scikit-image (v 0.20.0) package^40^. To calculate dendrite length, binary MAP2 channel images were skeletonized by scikit-image to 1 pixel representation whose length was measured by FilFinder (v 1.7.3) package^41^. For each cropped file, the puncta number divided by dendrite length produced the puncta density. All imaged regions from one batch of experiment were averaged to calculate the puncta density for one condition. 3 batches of data were plotted in total.
2. **Calcium imaging.** Time-lapsed videos of calcium imaging files were processed by CaImAn package^42^ to search for spiking somas and generate corresponding fluorescence intensity (ΔF/F) over time. The key parameters were: decay_time =0.4, p=1, gnb=2, merge_thr=0.85, rf=60, stride_cnmf=6, K=10, gSig=[40,40], method_init=‘greedy_roi’, ssub=1, tsub=1, min_SNR=200, rval_thr=0.85, cnn_thr=0.99, cnn_lowest=0.1. ΔF/F traces of all detected spiking somas from one field of view were averaged to produce one synchronized firing trace. SciPy (v1.10.1)^43^ algorithm “find_peaks” (height=0.15, width=(2,20), distance=20) was used to detect the spiking number and signal strength (ΔF/F) for each synchronized firing trace. The synchronizing firing rate was calculated by dividing spiking number against total time for each trace. To plot the firing rate (or ΔF/F) for each condition, the median of firing rate (or ΔF/F) from all traces of one batch was used. 3 batches of culture were plotted in total.
3. **Rabies tracing.** Coronal sections corresponding to Bregma -1.55 to -2.03 mm^44^ for hippocampal formation and Bregma -3.8 to -4.1 mm^44^ for lateral entorhinal cortex (LEC) were processed in Fiji by background subtraction. Regions of ipsilateral CA1, ipsilateral CA3, contralateral CA3, and ipisilateral LEC were cropped in Fiji for processing in scikit-image (v 0.20.0). Cropped regions were converted to binary images using the same threshold for the same channel, for the same batch of experiment. Binary regions containing more than 80 neighboring pixels (red channel for CA1) and 150 neighboring pixels (green channel for CA3 and LEC) were considered as neuron soma, and were counted by scikit-image^40^ functions “measure.label” and “measure.regionprops”. All counts from one mouse were used to calculate connectivity strength of ipsilateral CA3-CA1, contralateral CA3-CA1, LEC-CA1.
4. **cAMP imaging using Pink Flamindo 2**. After background subtraction, 488 and 546 channel signal from one field of view was used to calculate pink flamindo 2/GFP. 3-10 fields of view were imaged per condition per batch of culture. 3 batches of cultures were used in total.
5. **Phase-transitioned droplet.** After background subtraction, signals from indicated channels were used for analysis. 12.86µm linear region across the diameter of the droplets were used to plot the signal from the edge to the center of the droplets. To calculate the number of droplets per cluster, contacting droplet were counted as one cluster. scikit-image (v 0.20.0) package^40^ was used to count the size of droplets. 3 independent replicates were used for each experiment.

### Protein purification

6xHis-tagged GKAP, Shank3, Homer3 and PSD95 were purified as before^30^ with slight modifications. Constructs were transformed to BL21 (DE3) pLysS, which were induced at OD_600_=0.6 with 0.25mM IPTG at 16°C for 18hr. Cells were lysed in Ni-buffer A (20 mM Tris pH8, 500 mM NaCl, 5% glycerol, 4 mM BME, 20 mM Imidazole, 1x roche EDTA-free protease inhibitor, 100 U/ml benzonase) and cleared at SS34 rotor at 14000 rpm for 30 min at 4°C. Proteins were loaded onto Ni-NTA column, washed in Ni-buffer A and eluted in Ni-buffer B (20 mM Tris pH8, 250 mM NaCl, 5% glycerol, 4 mM BME, 400 mM Imidazole). His-tags were removed by 3C protease. Finally, the proteins were purified in size exclusion column (SD75 10/300 for GKAP, and SD200 10/300 for others) in SEC buffer (20 mM Tris pH8, 300 mM NaCl, 2 mM DTT). Lentiviruses containing CMV-TetO_2_-Flag-Lphn3 E31 and E32 were used to express in FreeStyle™ 293-F Cells^45^. 6xHis-tagged TEN2 and FLRT3 were cloned into pCMV vector and expressed in Expi293F^TM^ cells. 4 days post-transfection, medium were harvested and loaded onto Ni-NTA column, washed in Ni-buffer C (20 mM Hepes pH7.4, 150 mM NaCl, 20 mM Imidazole pH7.6) and eluted in Ni-buffer D (20 mM Hepes pH7.4, 150 mM NaCl, 250 mM Imidazole pH7.6). His-tags were removed by 3C protease, and the proteins were purified in size exclusion column (SD200 10/300) in SEC buffer (20 mM Hepes pH7.4, 150 mM NaCl).

### Fluorescence labeling of proteins

Proteins were labeled by NHS-dyes (AAT iFluor NHS-405, AAT iFluor-488, AAT iFluor NHS-546, Invitrogen Alexa NHS-647) were added to the protein at 1:1 molar ratio, and the labeling proceed at RT for 1hr. Reaction was quenched by 100 mM Tris pH8.2. Free dyes were removed by desalting column.

### Phase transition imaging and sedimentation assay

Protein mixtures were incubated at RT for 10 min. For imaging experiment, 5µl of sample was loaded onto channeled slide and imaged under Nikon confocal microscopy at 60x. For pelleting experiment, 10 µl of sample was centrifuged, and supernatant was immediately removed and pellet was resuspended in 2 x SDS loading buffer. All samples were subject to SDS-PAGE electrophoresis and stained in Coomassie G-250 blue.

### Statistical information

All numerical data were analyzed by two-sided Student’s t-test, except for the FPKM from RNAseq data which was analyzed by Wilconxon-rank sum test (*p < 0.05; **p < 0.01; ***p < 0.001). Non-significant results (p>0.05) are not identified. When applicable, boxplots (center line, median; box limits, first and third quartiles; minimal and maximum, minimal and maximum datapoint) show the distribution of datapoints (each point represents one datapoint).

## Extended Data Figures and Figure Legends

**Extended Data Figure 1:**
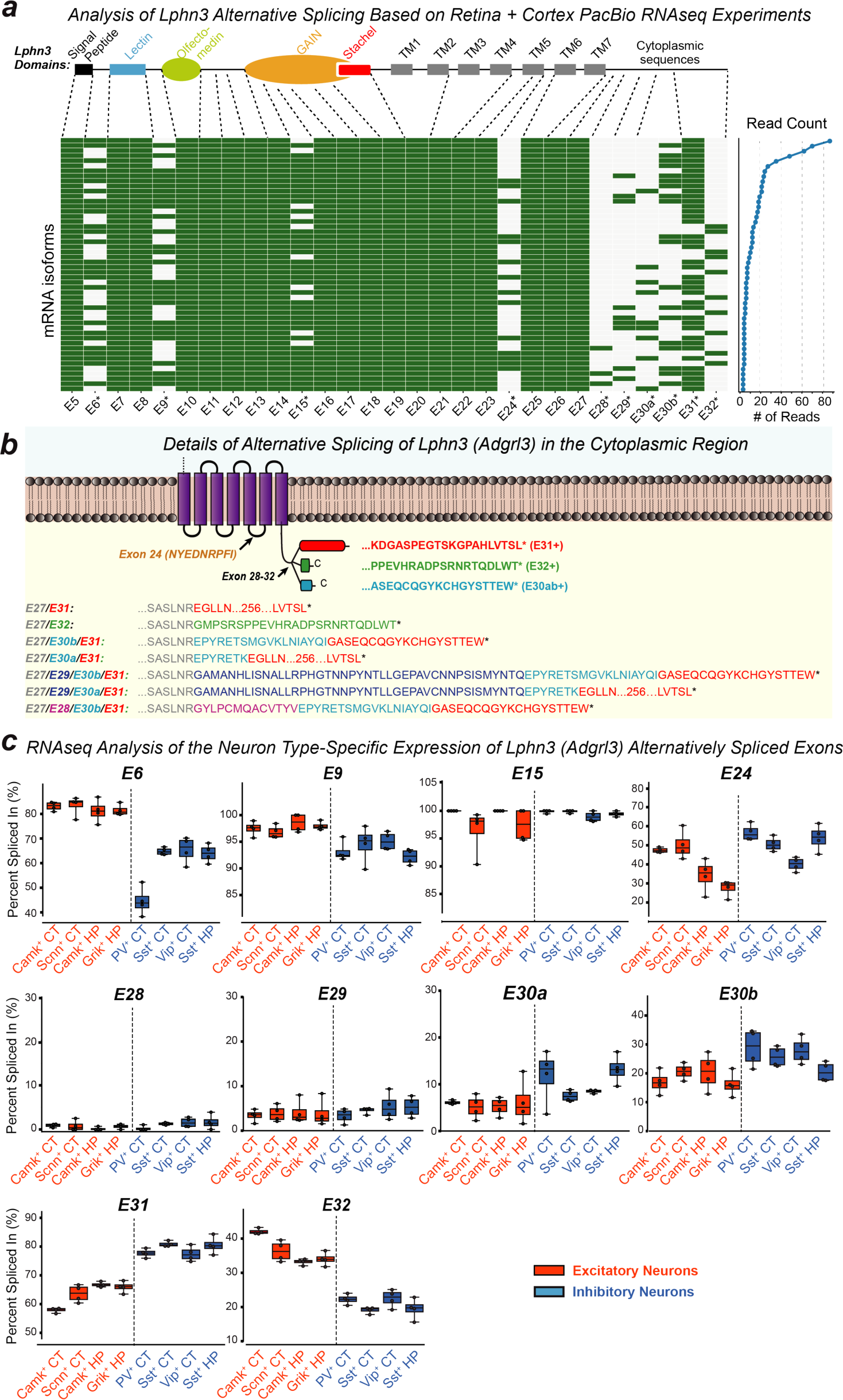
A subset of sites of alternative splicing of Lphn3 (*Adgrl3*) transcripts exhibits a high degree of cell-type specificity as revealed by RNAseq analyses. **a**, Analysis of a long-read PacBio sequencing data set from retina and cortex (NCBI BioProject repository; accession number PRJNA547800)^1^ uncovers extensive combinatorial alternative splicing of Lphn3 mRNAs. Reads are depicted as heatmaps (green boxes = Included exons; light gray boxes = excluded exons; exon numbers with asterisks = alternatively spliced exon). Each row represents a splice variant combination whose abundance is shown on the right (only the top 50 variants are shown). Note that for this figure and all following figures, ‘a’ and ‘b’ designations (such as ‘30a’ and ‘30b’) are alternative splicing donor or acceptor variants within an exon (see method “Generation of reference exon list” for details). For a similar analysis of Lphn2 and Lphn3, see ED Figure 2c, d. **b**, Detailed schematic of Lphn3 alternative splicing in the cytoplasmic region with the amino acid sequences of the resulting variants (asterisk = stop codon). Only the 7 TMR region and cytoplasmic sequences of Lphn3 are shown together with the major splice variants of the C-terminus. Note that whereas inclusion of the low-abundance exons E28, E29, and E30a introduces various possible variants, only three C-terminal Lphn3 sequence variants exist that are encoded by: (1) Exon 31 without the low-abundance Exon 30b, (2) Exon 31 with the low-abundance Exon 30b, and (3) Exon 32. **c**, Neuron type-specific alternative splicing of Lphn3. Raw data from a ribosome-associated transcriptome study^2^ were analyzed to calculate the abundance of each exon in PSI (percent spliced in) for 8 indicated neuron types from two brain regions (HP: hippocampus, CT: cortex). Each datapoint (n=4) represents one sample from 1 animal for excitatory neurons, and 2 animals for inhibitory neurons. The distribution of datapoints are shown in boxplots.

**Extended Data Figure 2:**
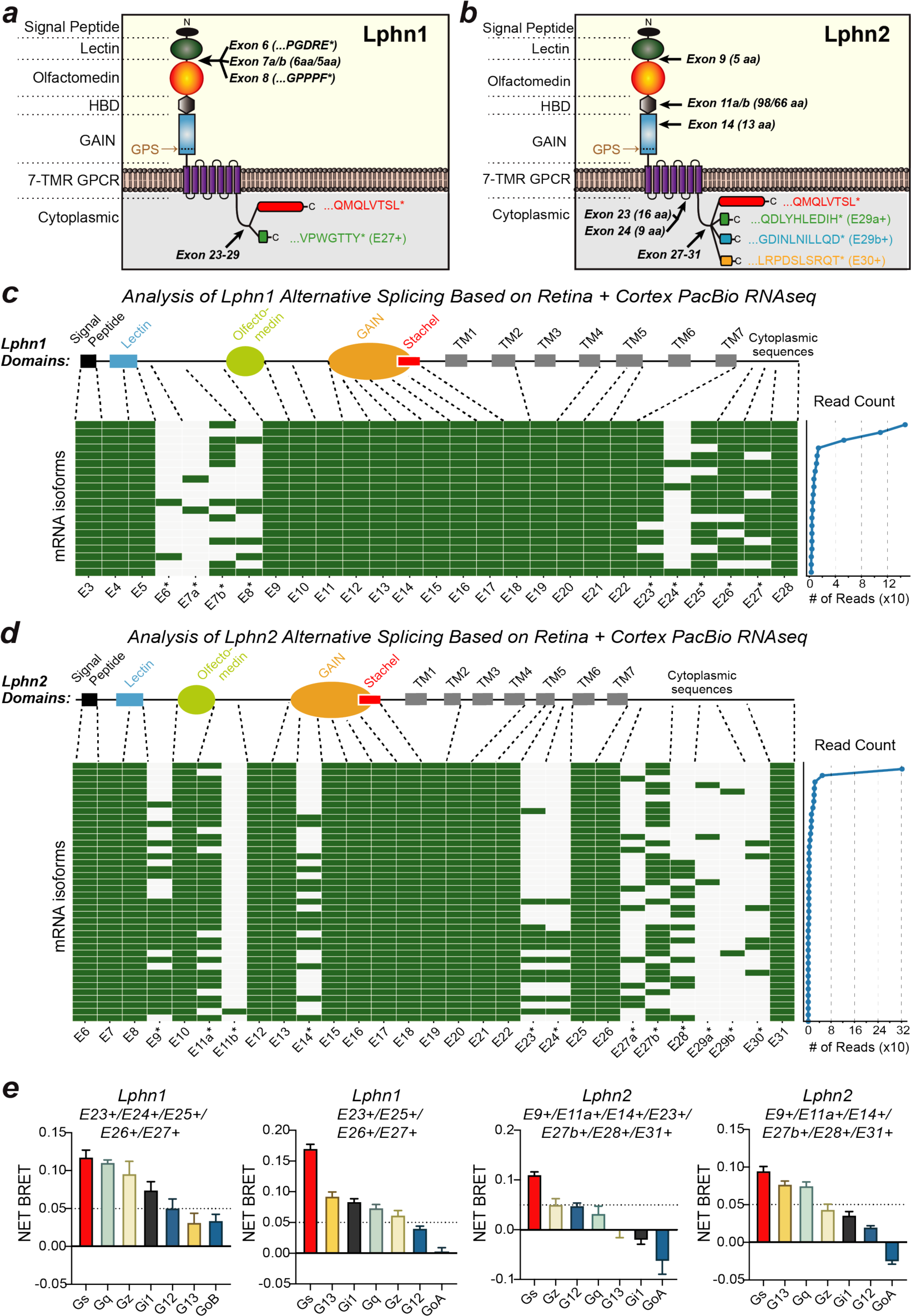
Regulation of G-protein coupling by alternative splicing of Lphn1 (*Adgrl1*) and Lphn2 (*Adgrl2*) **a** & **b**, Schematic of Lphn1 (a) and Lphn2 (b) alternative splicing with the amino acid sequences of some of the resulting variants (asterisk = stop codon). **c** & **d**, Analysis of a long-read PacBio sequencing data set from retina and cortex (NCBI BioProject repository; accession number PRJNA547800)^1^ also reveals extensive combinatorial alternative splicing of Lphn1 and Lphn2 mRNAs. Reads are depicted as heatmaps (green boxes = Included exons; light gray boxes = excluded exons; exon numbers with asterisks = alternatively spliced exon). Each row represents a splice variant combination whose abundance is shown on the right (only the most abundant variants are shown). **e**, G protein coupling preferences of two Lphn1 and Lphn2 splice variants revealed by TRUPATH analyses. The constitutive G-protein coupling strength (represented by the NET BRET signal) of the indicated Lphn1 and Lphn2 isoform were measured by TRUPATH assay in HEK293 cell. Splice variants with indicated spliced-in exons are shown. BRET signals at 300 ng receptor-transfected condition were normalized to the 0 ng transfected baseline. Graphs show means ± SEM from three batches of experiments (n=3).

**Extended Data Figure 3:**
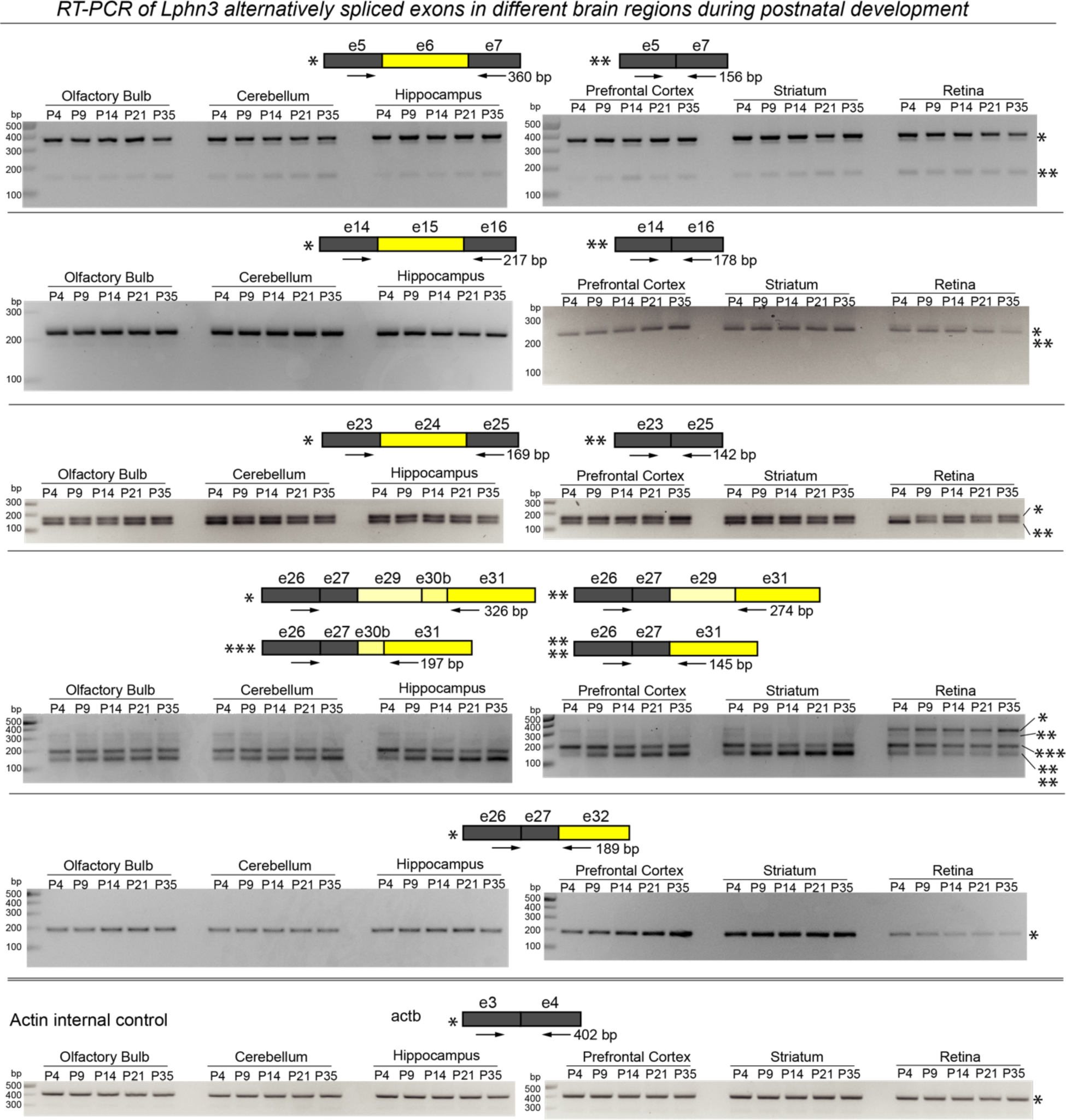
Pattern of Lphn3 alternative splicing analyzed by RT-PCR in different brain regions and at different times of postnatal development. Total RNA isolated from the indicated brain regions of C57BL/6 mice at the indicated postnatal developmental timepoints were analyzed by RT-PCRs using primers (labeled in arrows) at exon-exon junctions. RT-PCR products were separated by 2% agarose gel electrophoresis; bands are marked based on the predicted sizes of the alternatively spliced variants shown above each gel.

**Extended Data Figure 4:**
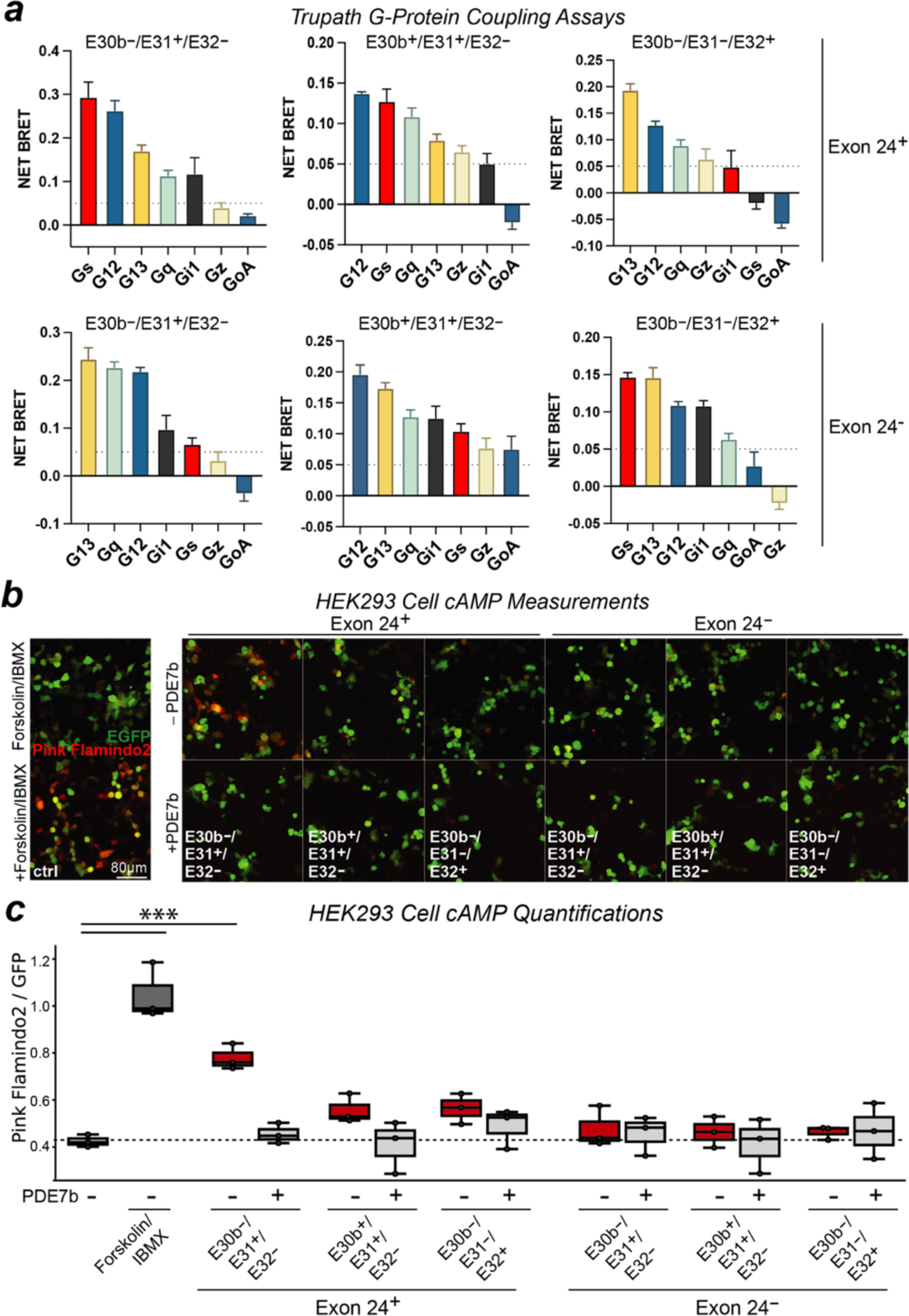
Detailed TRUPATH and Pink-flamindo2 reporter assay data for Lphn3 document splice variant-specific regulation of cAMP signaling by Lphn3. **a**, G protein coupling preference revealed by TRUPATH assays. The constitutive G-protein coupling strength (represented by NET BRET signal) of the indicated Lphn3 variants was measured using TRUPATH assays in HEK293 cell. BRET signals at the 300 ng receptor-transfected condition were normalized to 0 ng transfected baseline. Graphs show means ± SEM from three independent experiments (n=3). **b**, Representative images of cAMP assays. Intracellular cAMP levels were monitored in live HEK293T cells that had been transfected with Pink Flamindo2, EGFP and Gs(α/β/γ). Where indicated, Lphn3 variants and/or PDE7b (a cAMP-specific phosphodiesterase that blocks cAMP signaling) were co-transfected. Forskolin/IBMX were only applied to the control group. Imaging was performed by 10x confocal microscopy. **c**, Quantification of cAMP signals. The Pink Flamindo2 signal was normalized for the GFP signal in the same cells. Each datapoint represents one independent experiment (n = 3) as one batch of culture. Two-sided Student’s t-test was used to calculate the statistical significance (***p < 0.001).

**Extended Data Figure 5:**
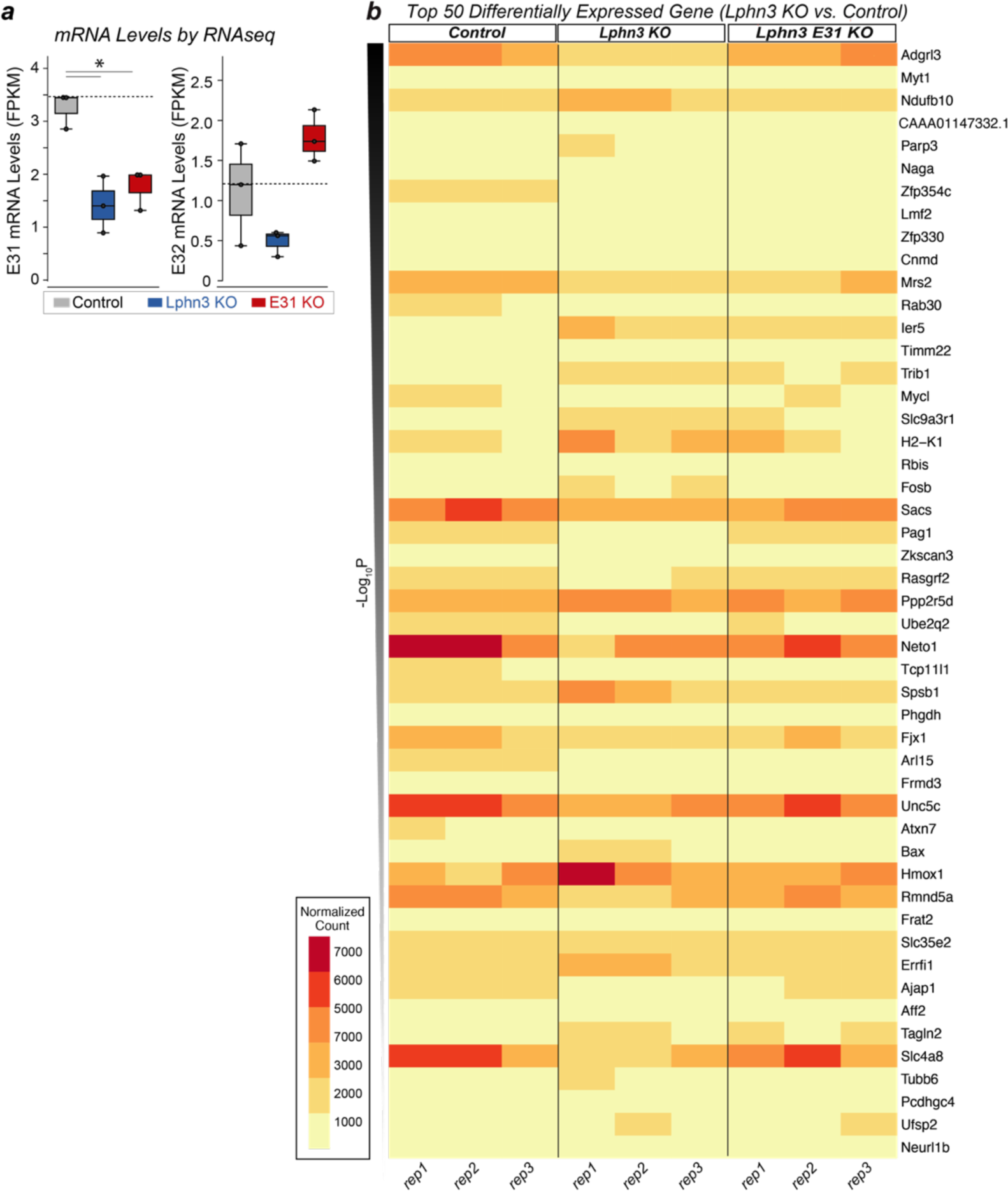
Transcriptomic level specificity of the Lphn3 KO and Lphn3 Exon31 KO. Transcriptomic data were obtained from hippocampal cultures that were infected with lentiviruses expressing control, Lphn3 KO, or Lphn3 Exon 31-specific KO gRNAs at DIV3 and analyzed at DIV14 by bulk RNAseq. **a**, Transcriptome quantifications by RNAseq confirm that the E31 KO and the Lphn3 KO similarly ablate expression of Exon 31-containing Lphn3 mRNAs (left) but have opposite effects on Exon 32- containing Lphn3 mRNAs (right). Each data point represents one independent culture (n=3); Wilconxon rank sum test was used to calculate the statistical significance (*p < 0.05). **b**, Heatmap illustrating the most significant changes in gene expression observed in three independent RNAseq experiments (rep1-rep3). Normalized count of each replicate for each condition is shown for the top 50 genes which are differentially expressed (based on the p value of Lphn3 KO vs control).

**Extended Data Figure 6:**
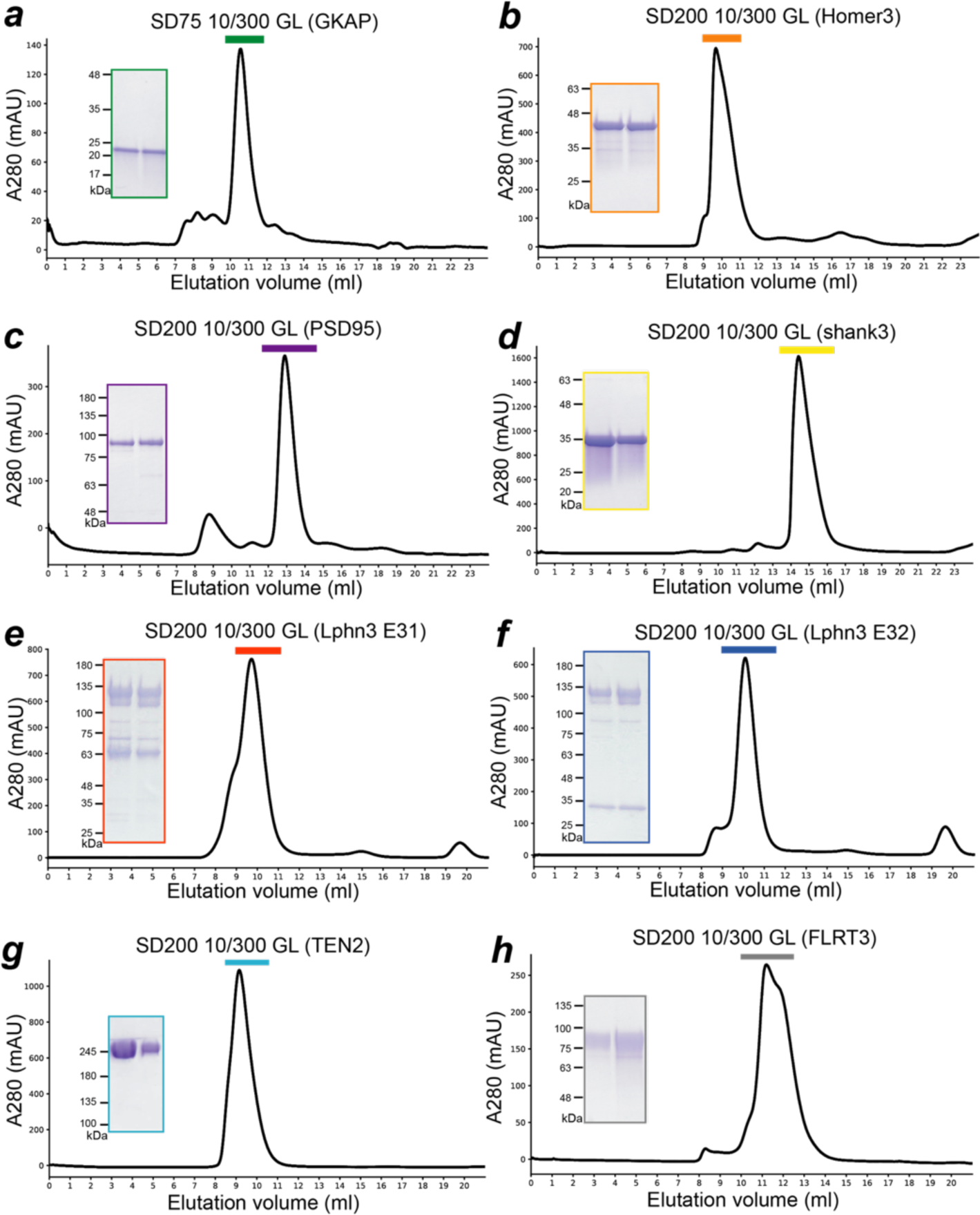
Quality control of purified proteins used for phase-transition experiments. **a-h**, Chromatographs of proteins on gel-filtration columns during the last step of purification, and analysis by SDS-PAGE and Coomassie staining of the purified proteins (insets). Peak regions (labeled bars) were pooled as the final purified samples for this study. SDS-PAGE was employed to analyze the purity of fractionated samples. Note that Lphn3 is autocleaved in the GAIN domain to produce amino-terminal (apparent molecular weight ∼130kD) and carboxyl-terminal fragments (apparent molecular weight ∼65kD for Lphn3 E31 and ∼30kD for Lphn3 E32) which are non-covalently connected within the GAIN domain^3^.

**Extended Data Figure 7:**
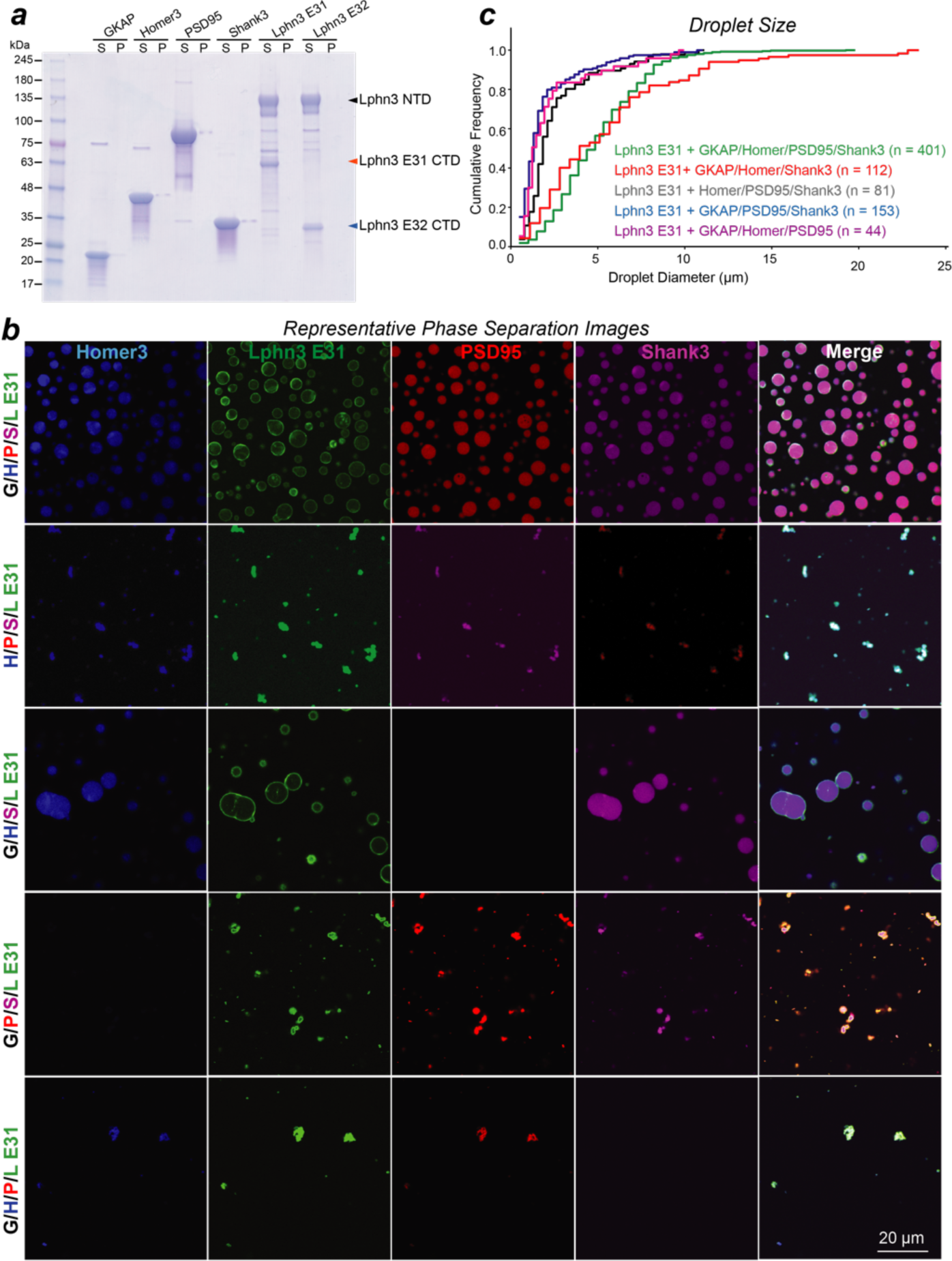
Further characterization of the Lphn3 association with postsynaptic scaffolds undergoing phase separation. **a**, Sedimentation behavior of individual proteins. Purified proteins were diluted to the same concentration, and centrifuged in the same condition, as used in the phase-transition assay. Supernatant (S) and pellet (P) were subject to SDS-PAGE for analysis. **b**, Contribution of individual postsynaptic scaffold protein to phase transitions in the presence of Lphn3 containing Exon 31. Individual scaffold proteins were omitted during the phase separation and droplets were subjected to the same imaging condition at 60x. Homer3, Lphn3, PSD95, Shank3 were labeled by NHS-ester fluorophore 405, 488, 546, 647, respectively. G: GKAP, H: Homer3, P: PSD95, S: Shank3, L E31: Lphn3 E31. Note that only PSD95 is dispensable for formation of large droplets. **c**, Quantification of phase-transitioned droplet diameter under the conditions shown in **b**.

**Extended Data Figure 8:**
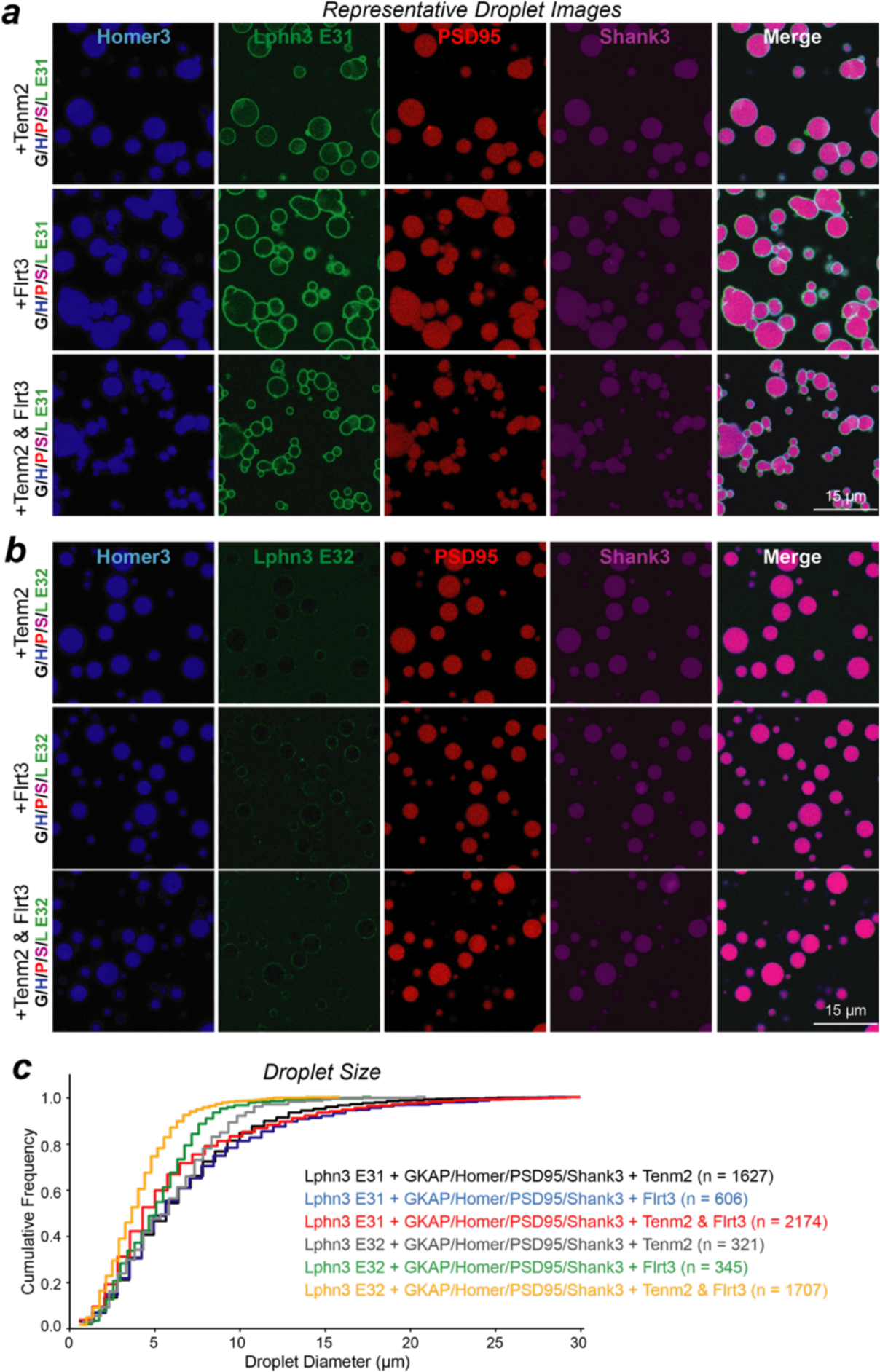
Further analyses of the effect of presynaptic Lphn3 ligands on the clustering of phase transition droplets formed by postsynaptic scaffolds with Lphn3 E31. **a**, Imaging of postsynaptic scaffold proteins (GHPS) with Lphn3 E31, containing 10 µM presynaptic ligands Tenm2 or/and Flrt3. **b**, Imaging of postsynaptic scaffold proteins (GHPS) with Lphn3 E32, containing 10 µM presynaptic ligands Tenm2 or/and Flrt3. **c**, Quantification of phase-transitioned droplet diameter under the conditions shown in **a** and **b**. Note that although only Lphn3 E31 but not Lphn3 E32 associates with phase-transitioned droplets, Lphn3 E31 has no major effect on droplet size.

